# A developmentally regulated long-range enhancer-promoter contact mediates human neural development

**DOI:** 10.64898/2025.12.01.691702

**Authors:** Devin Bready, Shuai Wang, Niklas Ravn-Boess, Joshua Frenster, Jonathan Sabio, Robert Kushmakov, Finnegan Clark, Adler Guerrero, Hannah Dada, Cathryn Lapierre, Kristyn Galbraith, Catherine Do, Priscillia Lhoumaud, Jod Prado, Albert Jiang, Sara Haddock, Claire D. Kim, Matija Snuderl, Timothée Lionnet, Aristotelis Tsirigos, Jane Skok, Dimitris G. Placantonakis

**Affiliations:** Department of Neurosurgery, New York University Grossman School of Medicine, NY; Medical Scientist Training Program, New York University Grossman School of Medicine, NY; Institute for Systems Genetics, New York University Grossman School of Medicine, NY; Department of Pathology, New York University Grossman School of Medicine, NY; Perlmutter Cancer Center, NYU Langone Health, NY; Department of Cell Biology, New York University Grossman School of Medicine, NY; Department of Medicine, New York University Grossman School of Medicine, NY; Applied Bioinformatics Laboratory, New York University Grossman School of Medicine, NY; Neuroscience Institute, New York University Grossman School of Medicine, NY

**Keywords:** *SOX2*, enhancer, CTCF, chromatin organization, neuroectoderm, forebrain, gene regulation, development

## Abstract

SOX2 is a core pluripotency factor in human embryonic stem cells (hESCs), but upon differentiation to the three germ layers, its expression is preserved selectively in neuroectoderm. The mechanisms regulating *SOX2* transcription in distinct developmental stages remain incompletely understood. Here, we demonstrate that a distant enhancer 550 kb from the human *SOX2* locus is selectively activated in neural stem cells (NSCs) and establishes long-range contact with the *SOX2* gene. CRISPR-Cas9 excision of the enhancer has no effect in hESCs but reduces *SOX2* transcription in NSCs and impairs neuroectodermal differentiation and forebrain specification in teratomas and cerebral organoids. CRISPR excision of a CTCF recognition motif adjacent to the enhancer does not affect enhancer activation in neuroectoderm but reduces chromatin looping and *SOX2* transcription to partially reproduce phenotypes seen with enhancer deletion. Our findings indicate that the development of the human nervous system depends on a developmentally regulated long-range contact between a distant enhancer and the *SOX2* locus.

## INTRODUCTION

Embryonic development relies on spatiotemporally orchestrated lineage commitment and differentiation steps that start with pluripotent stem cells (PSCs) in the inner cell mass and lead to the full gamut of cell fates that populate the embryo. Each one of these steps requires lineage-defining transcriptional programs, which are determined by specific transcription factors (TFs) and epigenetic mechanisms^1,2^ that include both coding and non-coding regions of the genome. Within the non-coding genome, enhancers are regulatory elements that can influence transcription at target loci at a distance through interactions mediated by three-dimensional (3D) chromatin folding. One tier of 3D chromatin organization involves topologically associating domains (TADs), which are 0.5-1.0 Mb-long chromatin regions that facilitate intra-TAD 3D contacts^3,4^ and whose formation relies on the cohesin complex and DNA-binding chromatin organizer CTCF.^5,6^ The dynamic regulation of TAD organization and 3D chromatin contacts in normal development and disease is an area of active investigation.^7,8^

SOX2 belongs to the SOX (SRY-related high-mobility group box) family of TFs, which includes 20 separate SOX proteins classified into 9 subfamilies.^9^ Within the SOXB1 subfamily, SOX2 serves as a core pluripotency TF alongside OCT4 and NANOG in embryonic stem cells (ESCs).^10,11,12^ These core pluripotency TFs support transcription of each other by forming heteromultimers in a positive-feedback mechanism,^13,14^ and define a global transcriptome favoring pluripotency and preventing differentiation. Interestingly, SOX2 retains an important functional role after exit from the ESC stage. In embryogenesis, the specification of the three germ layers requires that SOX2 persist only in neuroectodermal progenitors, where it is essential for neural development.^15,16^ In contrast, *SOX2* transcription is silenced in early mesodermal and endodermal lineages, with subsequent expression in endodermal lineages of the developing lung, trachea, stomach, and esophagus.^16–18^ In the neural tube, SOX2 is ubiquitously expressed in the absence of other pluripotency TFs, and is required for the self-renewal and multipotency of neural progenitors.^19–25^ It partners with other lineage-specific transcription factors, such as PAX6, to transactivate relevant sets of genes that define neural stem cells (NSCs),^26,27^ and helps maintain bivalent chromatin marks on neuronal differentiation genes, thereby keeping them poised and readily activated when neuronal differentiation programs are initiated.^19–24^ This raises the question of how *SOX2* transcription is regulated in neural progenitors relative to ESCs, given the absence of other pluripotency factors in the neural lineage. Furthermore, SOX2 is also expressed and functionally relevant in tumors deriving from neural tissue, such as glioma.^28–30^ These findings underscore the importance of characterizing transcriptional regulatory mechanisms for SOX2, a gene critical to pluripotency, central nervous system development and tumorigenesis.

Previous work has characterized enhancer elements that regulate *SOX2* transcription in the mammalian nervous system, primarily using mouse models.^31–36^ However, the organization of the genome around the *SOX2* locus in human central nervous system development remains incompletely understood. Here we show that, in human neural progenitors, *SOX2* transcription depends on a long-range contact with a distant enhancer located ∼550 kb away within the same TAD. Activation of the enhancer and establishment of the long-range contact with the *SOX2* locus are developmentally regulated and established selectively in neural progenitors but not in human ESCs (hESCs). CRISPR-mediated excision of the enhancer does not affect hESC self-renewal, but results in impaired neural development and neurogenesis, particularly regarding the generation of forebrain lineages. Our study complements previous work on the regulation of *SOX2* transcription, but also demonstrates a developmentally regulated distant enhancer and 3D chromatin loop crucial to neural and forebrain development.^33,37,38^ This mechanism highlights the impact of dynamic reconfiguration of the non-coding genome and 3D organization of chromatin during development.

## RESULTS

### A phylogenetically conserved and neuroectoderm-specific long-range contact between a distant enhancer and the human SOX2 locus

Our previous work suggested that, in hESC-derived NSCs, the *SOX2* gene locus in chromosome 3q makes a strong long-range contact with a region ∼550 kb telomeric to it within the canonical TAD of this genomic neighborhood in 4C-seq assays^39^ (Figure 1A). This region is marked by a CTCF cognate motif (numbered 5 and termed *nMotif* hereafter), while the *SOX2* locus is also marked by several CTCF motifs (numbered 1-4) (Figure 1A). Our Assay for Transposase-Accessible Chromatin sequencing (ATAC-seq) analysis indicated that, immediately centromeric (∼16 kb) to *nMotif* lies a ∼5.6 kb genomic region with the most accessible chromatin in the TAD besides the *SOX2* locus (Figure 1A). In addition, chromatin immunoprecipitation sequencing (ChIP-seq) showed that this region is highly decorated with the euchromatic histone modification H3K27ac (acetylation) but lacks the facultative repressive modification H3K27me3 (trimethylation) (Figure 1A), suggesting it may represent a transcriptionally active enhancer element. Analysis with HiC and virtual 4C validated this chromatin loop in NSCs (Figure 1B), and ChIP-seq for CTCF demonstrated CTCF occupancy at *nMotif*, as well as around the *SOX2* gene locus (Figure 1B) in NSCs.

**Figure 1.**
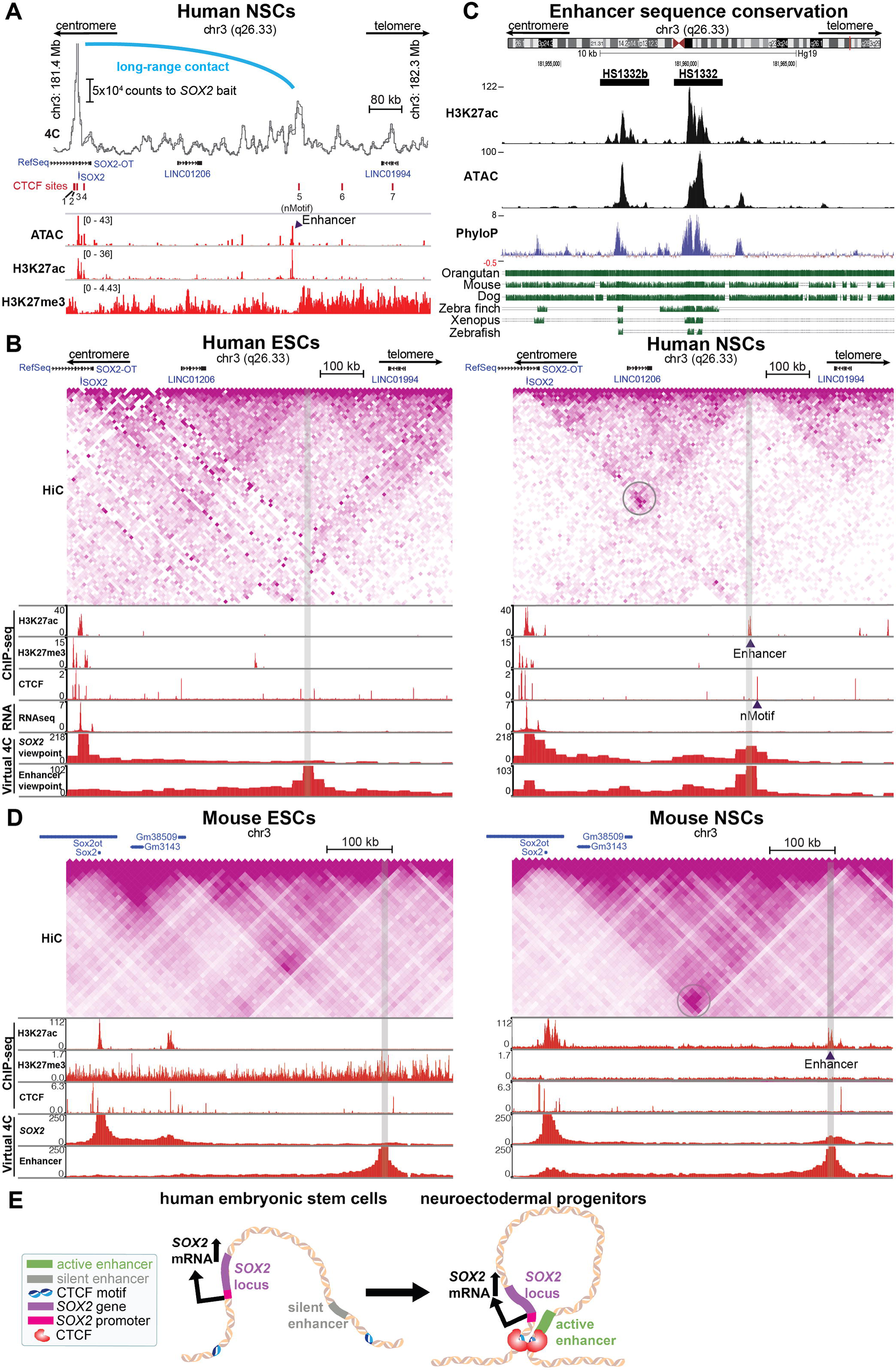
A conserved and neuroectoderm-specific long-range contact between a distant enhancer and the *SOX2* locus. (A) Genomic organization of a 1 Mb domain of the *SOX2* TAD in human NSCs. 4C-sequencing with a *SOX2* bait, ATAC-Sequencing, and H3K27ac and H3K27me3 ChIP-sequencing are shown. (B) Comparison of hESC and NSC HiC sequencing, ChIP-sequencing for CTCF, H3K27ac, and H3K27me3, RNA-sequencing transcript reads, and virtual 4C-Sequencing visualizations (derived from our HiC sequencing) with a viewpoint of either the *SOX2* gene promoter or the putative enhancer region. See also Figure S1A,B. (C) Genome browser view of *HS1332* and adjacent *HS1332b* sequences corresponding to the putative enhancer in the human genome with H3K27ac ChIP-sequencing and ATAC-sequencing from NSCs, PhyloP score for 100 vertebrate genomes, and view of conserved sequences (green bars) for 6 select species. See also Figure S1C. (D) Comparison of mouse ESC and NSC HiC sequencing, ChIP-sequencing for CTCF, H3K27ac, and H3K27me3, and virtual 4C-Sequencing visualizations (derived from HiC sequencing, GSE96107) with a viewpoint of either the *SOX2* gene promoter or the putative enhancer region. (E) Schematic demonstrating the hypothesis tested.

We then tested whether this putative enhancer-promoter interaction may be relevant in other cell lineages. We interrogated data from the Epigenome Roadmap in hESCs,^40^ which also express *SOX2*, and mesodermal progenitors,^41^ which do not. While both NSCs and hESCs robustly transcribe *SOX2*, the enhancer has an open chromatin conformation on ATAC-seq and is decorated with the euchromatic modification H3K27ac in ChIP-seq analysis only in NSCs, but not hESCs (Figure 1B, Figure S1A). Similarly, analysis of HiC data with virtual 4C, utilizing either the *SOX2* genomic locus or the distant enhancer as the viewpoint, indicated that the long-range contact is significantly more prominent in NSCs than hESCs (Figure 1B, Figure S1B). We found no evidence for enhancer activation or chromatin looping in public domain ChIP-seq and HiC from mesodermal progenitors,^41^ which do not express *SOX2* (Figure S1A,B). These data suggested the developmental co-regulation of this enhancer and the associated chromatin loop upon neuralization.

Within the genomic region corresponding to this putative enhancer lie two H3K27ac peaks, one of which was previously designated *HS1332* in the VISTA Enhancer dataset^42^ (we will refer to the other peak as *HS1332b* hereafter). Further analysis suggested that the genomic sequence of this enhancer element, particularly *HS1332*, is phylogenetically conserved from zebrafish to humans (Figure 1C). Strikingly, when the vertebrate conservation scores are aligned against ChIP-Seq and ATAC-Seq of NSCs, the loci displaying the highest amount of H3K27ac are those corresponding to the conserved regions (Figure 1C). Of note, the nMotif sequence overlaps with element *mm639* in the VISTA dataset, which did not demonstrate any enhancer activity by itself.

The evolutionary conservation of the enhancer sequence itself led us to ask if the same phenomenon of TAD reorganization and putative enhancer activation^42^ could be seen during neuralization in other species. In mice, the analogous enhancer is approximately 450 kb telomeric to *SOX2* and displays a robust degree of conservation^37^ (Figure 1C,D). As in humans, while the long-range contact was absent and the enhancer was silent in mouse ESCs, upon neuralization the long-range contact surrounding the enhancer and the activation of the distal enhancer itself were robust (Figure 1D), consistent with previous observations^37,43–45^. While relevant datasets in zebrafish were sparse, we found evidence that the evolutionarily conserved enhancer element, which in the zebrafish genome is approximately 45 kb distal to *SOX2*, is also activated and recruited in embryonic brain tissue but not muscle (Figure S1C).^46^ Collectively, these data lead us to hypothesize that this conserved enhancer element acts on the *SOX2* gene locus via a CTCF-mediated long-range chromatin loop to support *SOX2* transcription specifically in embryonic neural development but not in the pluripotent state (Figure 1E).

### Excision of the distant enhancer or disruption of the chromatin loop impair specification of neuroectodermal lineages

To mechanistically investigate the role the long-range enhancer-*SOX2* contact plays in human development, we used a dual gRNA CRISPR-Cas9 approach to excise the 2.7 kb *HS1332* enhancer element (Δ*HS1332*) in H9 hESCs transgenic for the *Hes5::GFP* bacterial artificial chromosome (BAC)^47–49^ (Figure 2A). This transgene allows identification of neural progenitors through expression of GFP. We implemented the same approach to excise the CTCF motif (90 bp) adjacent to the enhancer (Δ*nMotif*) (Figure 2A). A single gRNA targeting the human *ROSA26* locus was used as control. hESCs were transduced with lentiviruses expressing Cas9 and each respective gRNA combination and pools of cells were isolated by FACS. To generate hESC clones with the desired genetic alterations, the FACS-isolated hESCs were plated on mouse embryonic fibroblasts (MEFs) in 24-well tissue culture plates at a nominal seeding density of 5 cells/well, which was experimentally determined to yield one clone per 24 wells seeded. Deletions in each condition were genotyped with PCR and Sanger sequencing (Figure 2B, Figure S2A). In anticipation of a lower efficiency in genome editing when excising the large *HS1332* element, further subcloning was performed. Among five further subclones generated, one was found to have the desired deletion by PCR and Sanger sequencing (Figure 2B, Figure S2A). We also attempted to excise both *HS1132* and *HS1332b* elements as a single cassette but were not successful.

**Figure 2.**
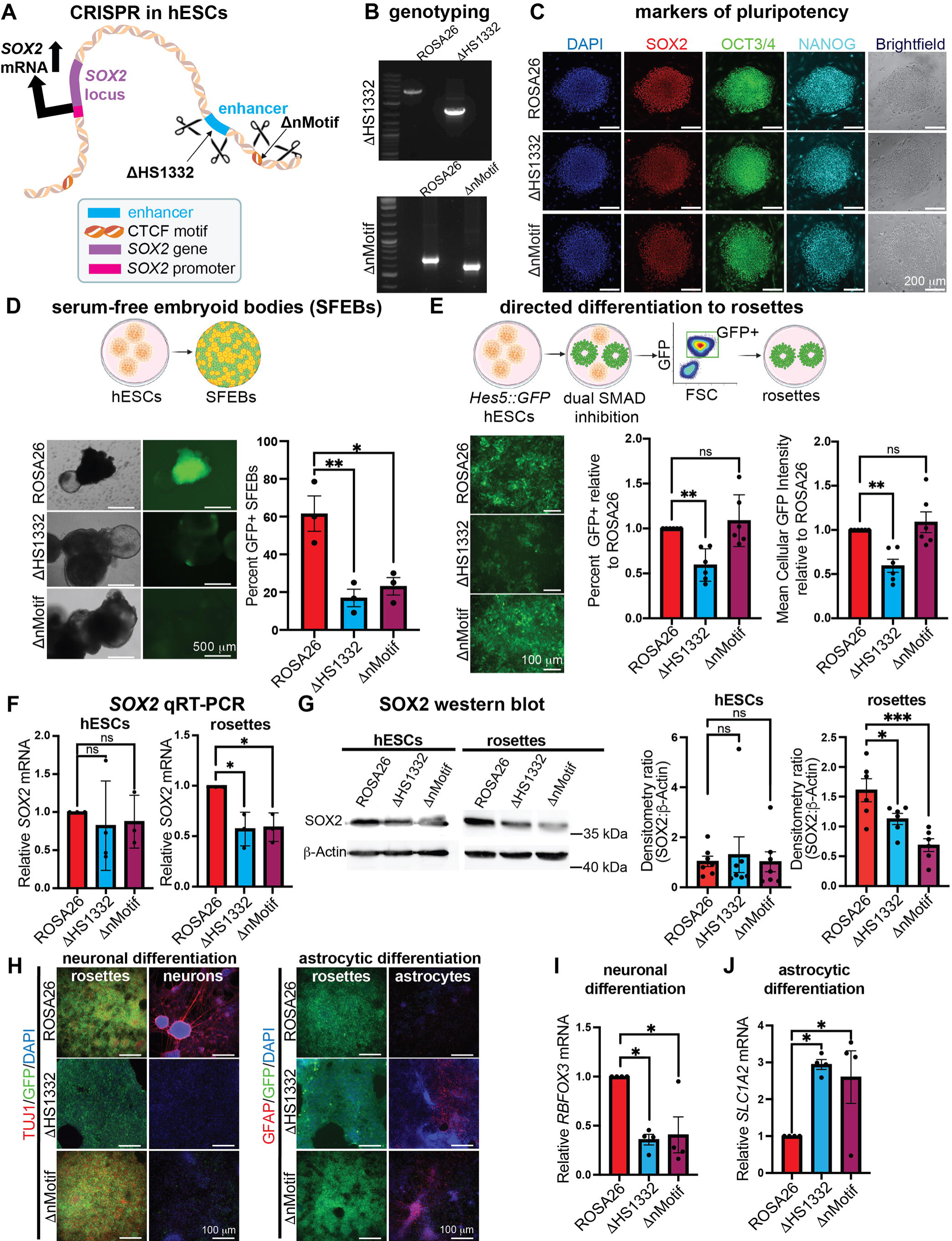
Disruption of *HS1332* and the associated 3D chromatin loop impair neuralization, SOX2 expression and neuronal differentiation. (A) Graphical representation of targeted deletion of either the *HS1332* enhancer or the CTCF *nMotif*. (B) Gel electrophoresis of PCR products of the *HS1332*-containing domain in *ROSA26* vs Δ*HS1332* hESCs (top), or the *nMotif*-containing domain in *ROSA26* vs Δ*nMotif* hESCs (bottom). Additional information is found in Figures S2, S3. (C) Representative immunofluorescence images for hESC markers SOX2 (red), OCT4 (green), Nanog (teal) across the *ROSA26*, Δ*HS1332*, and Δ*nMotif* conditions at the hESC stage. Nuclei were counterstained with DAPI. (D) Graphical representation of embryoid body (SFEB) formation experiments highlighting the emergence of GFP expression from the *Hes5::GFP* reporter (top). Representative images from *ROSA26*, Δ*HS1332*, and Δn*Motif* SFEBs (bottom left). The bar graph (bottom right) represents the quantification of GFP-positive SFEBs (n=3 biological replicates, one-way ANOVA F_2,6_=13.43, p <0.01; Tukey’s *post hoc* multiple comparison test: *ROSA26* vs Δ*HS1332* **p<0.01; *ROSA26* vs Δ*nMotif* *p<0.05). (E) Graphical representation of directed *in vitro* differentiation to neural rosettes (top) and representative wide field fluorescence images of each line (bottom left). The bar graph in the middle shows quantification of the percentage of GFP-positive (GFP+) cells after directed differentiation as measured via flow cytometry, normalized to the *ROSA26* condition in each biological replicate (n=6 biological replicates, one-way ANOVA F_2,15_=10.76, p=0.0013; Tukey’s *post hoc* multiple comparison test: *ROSA26* vs Δ*HS1332* **p<0.01). ns, not significant. The bar graph on the right shows mean GFP intensity in rosettes across experimental conditions (n=6 biological replicates, one-way ANOVA F_2,15_=5.921, p=0.0228; Tukey’s *post hoc* multiple comparison test: *ROSA26* vs Δ*HS1332* **p<0.01). ns, not significant. (F) qRT-PCR analysis shows no change in *SOX2* transcript levels at the hESC stage (n = 3 biological replicates, one-way ANOVA F_2,7_=0.1535, p=0.8605) and a decrease in *SOX2* transcript in Δ*HS1332* and Δn*Motif* rosettes (n=4 biological replicates, ANOVA F_2,6_=11.11, p=0.0096; Tukey’s *post hoc* multiple comparison test: *ROSA26* vs Δ*HS1332* *p < 0.05, *ROSA26* vs Δ*nMotif* *p < 0.05). ns, not significant. (G) Representative immunoblots for SOX2 and β-Actin show SOX2 downregulation in rosettes but not hESCs in the Δ*HS1332* and Δn*Motif* conditions (left). Bar graphs (right) demonstrate summary densitometry statistics for hESCs (n=7 biological replicates, one-way ANOVA F_2,18_=0.1062, p=0.8998; Tukey’s *post hoc* multiple comparison test: *ROSA26* vs Δ*HS1332* p>0.05; *ROSA26* vs Δ*nMotif* p>0.05) and rosettes (n=6 biological replicates, one-way ANOVA F_2,15_=11.10, p=0.0011; Tukey’s *post hoc* multiple comparison test: *ROSA26* vs Δ*HS1332* *p<0.05, *ROSA26* vs Δ*nMotif* ***p<0.001). ns, not significant. (H) Immunofluorescence microscopy before and after differentiation of rosettes to neurons (left) and astrocytes (right). (I) qRT-PCR for *RBFOX3* transcript normalized to *GAPDH* following 2 weeks of neuronal differentiation with BDNF and ascorbic acid (n=4 biological replicates, one-way ANOVA F_2,8_=8.999, p<0.01; Tukey’s *post hoc* multiple comparison test: *ROSA26* vs Δ*HS1332* *p < 0.05, *ROSA26* vs Δ*nMotif* *p < 0.05). (J) qRT-PCR for *SLC1A2* transcript normalized to *GAPDH* following 2 weeks of astrocytic differentiation with 4% FBS (n=4 biological replicates, one-way ANOVA F_2,8_=7.469, p=0.0104; Tukey’s *post hoc* multiple comparison test: *ROSA26* vs Δ*HS1332* *p<0.05, *ROSA26* vs Δ*nMotif* *p<0.05).

One clone from each of the Δ*HS1332* and Δ*nMotif* conditions was chosen for further experimentation. To confirm the deletions, we performed bulk whole genome sequencing (WGS) of hESCs from the Δ*HS1332* and Δ*nMotif* clones (Figure S2B,C), which demonstrated homozygous deletion of *nMotif* but deletion of only one of the two *HS1332* alleles. To further investigate the Δ*HS1332* condition, we generated a panel of fluorescence *in situ* hybridization (FISH) probes against the 2.7 kb deleted region of the *HS1332* enhancer and a nearby 45 kb region (∼300 kb telomeric), which was not targeted for deletion (Figure S3A). A universal anchor control probe set targeting a 20 kb locus near the *HBB* (hemoglobin subunit beta) gene on chromosome 11, with known labeling efficacy, was included as an additional control. The *HS1332* probe set was expected to have low labeling efficiency given the small size of the genomic region interrogated (2.7kb). While ∼70% of *ROSA26* hESCs and hESC-derived rosette neuroectodermal stem cell (rosette) cultures, which developmentally resemble neural plate precursors,^50,51^ manifested *HS1332* FISH labeling, only ∼30-40% of their Δ*HS1332* counterparts showed *HS1332* labeling (Figure S3B-E). Collectively, these data suggested homozygous deletion of Δ*nMotif* and hemizygous deletion of *HS1332*. A less likely possibility is that the Δ*HS1332* condition represents a mix of cells with homozygous *HS1332* deletion and wild-type cells. Finally, we utilized WGS data to generate pseudo-karyotypes that demonstrated no significant differences in chromosomal integrity among the Δ*HS1332*, Δ*nMotif* and *ROSA26* conditions (Figure S4).

Hemizygous excision of the enhancer or homozygous deletion of the nMotif had no impact on the self-renewal of hESCs, as evidenced by expression of pluripotency transcription factors (OCT4, NANOG, SOX2) and cell surface markers (SSEA4, TRA-1-81) (Figure 2C, Figure S5A,B). To assess the effects of these modifications in neural differentiation, we performed two types of assays. First, we employed *in vitro* unbiased differentiation of Δ*HS1332* or Δ*nMotif* hESCs as serum-free embryoid bodies (SFEBs).^48,49^ These assays indicated significant impairment in generation of GFP+ neuroectodermal precursors derived from Δ*HS1332* or Δ*nMotif* hESCs relative to the *ROSA26* control (Figure 2D). Second, we used dual SMAD inhibition to direct differentiation of hESCs to neural rosettes.^50,51^ These assays also demonstrated significant decreases in both the number of GFP+ progenitors and the intensity of GFP fluorescence in the Δ*HS1332* condition, but no significant effect in the Δ*nMotif* condition (Figure 2E). These findings suggested that perturbing the enhancer and long-range chromatin loop had no effect on hESC self-renewal but impaired specification of neuro-ectoderm.

In both excision groups, impaired neural specification correlated with significant reductions in *SOX2* transcript levels in rosettes, while no differences were observed at the hESC stage (Figure 2F). Immunoblotting for SOX2 confirmed that SOX2 protein levels were reduced at the rosette stage in the Δ*HS1332* and Δ*nMotif* conditions, with no differences seen at the hESC stage (Figure 2G). Directed neuroglial differentiation of the *SOX2*-deficient Δ*HS1332* and Δ*nMotif* rosette progenitors reproducibly failed to generate neurons, while generation of astrocytes was facilitated, as evidenced by immunostaining for beta-tubulin III (TUJ1; neuronal) and glial fibrillary acidic protein (GFAP; astrocytic), and qRT-PCR for *RBFOX3* (neuronal) and *SLC1A2* (astrocytic) mRNA (Figure 2H-J). Collectively, these data suggest that perturbation of the long-range enhancer-*SOX2* locus contact interferes with the development of neuroectodermal progenitors and subsequent neuroglial differentiation.

To validate these *in vitro* findings in an *in vivo* setting, we performed teratoma assays in NOD.Cg- *Prkdc^scid^ Il2rγ^tm1Wjl^*/SzJ (NSG) mice by implanting *ROSA26,* Δ*HS1332*, and Δ*nMotif* hESCs in the flank. Teratomas can serve as a model for studying development because they recapitulate a large number of diverse cell types, including distinct neural lineages.^52^ After allowing teratomas to form for six weeks (Figure 3A), we extracted them and performed blinded histologic analysis of hematoxylin and eosin (H&E) sections. This assay indicated impairment in neuroectodermal rosette formation in Δ*HS1332* and Δ*nMotif* teratomas (Figure S6A,B). Immunofluorescence microscopy of the teratomas revealed a significantly reduced prevalence of nuclei positive for SOX2 and the neuroectodermal TF PAX6 in the excision groups. Interestingly, Δ*HS1332* teratomas, and to a lesser extent Δ*nMotif* teratomas, showed enrichment for the endodermal TF HNF3B, while no differences were found for the mesodermal marker smooth muscle actin (SMA) (Figure 3B,C).

**Figure 3.**
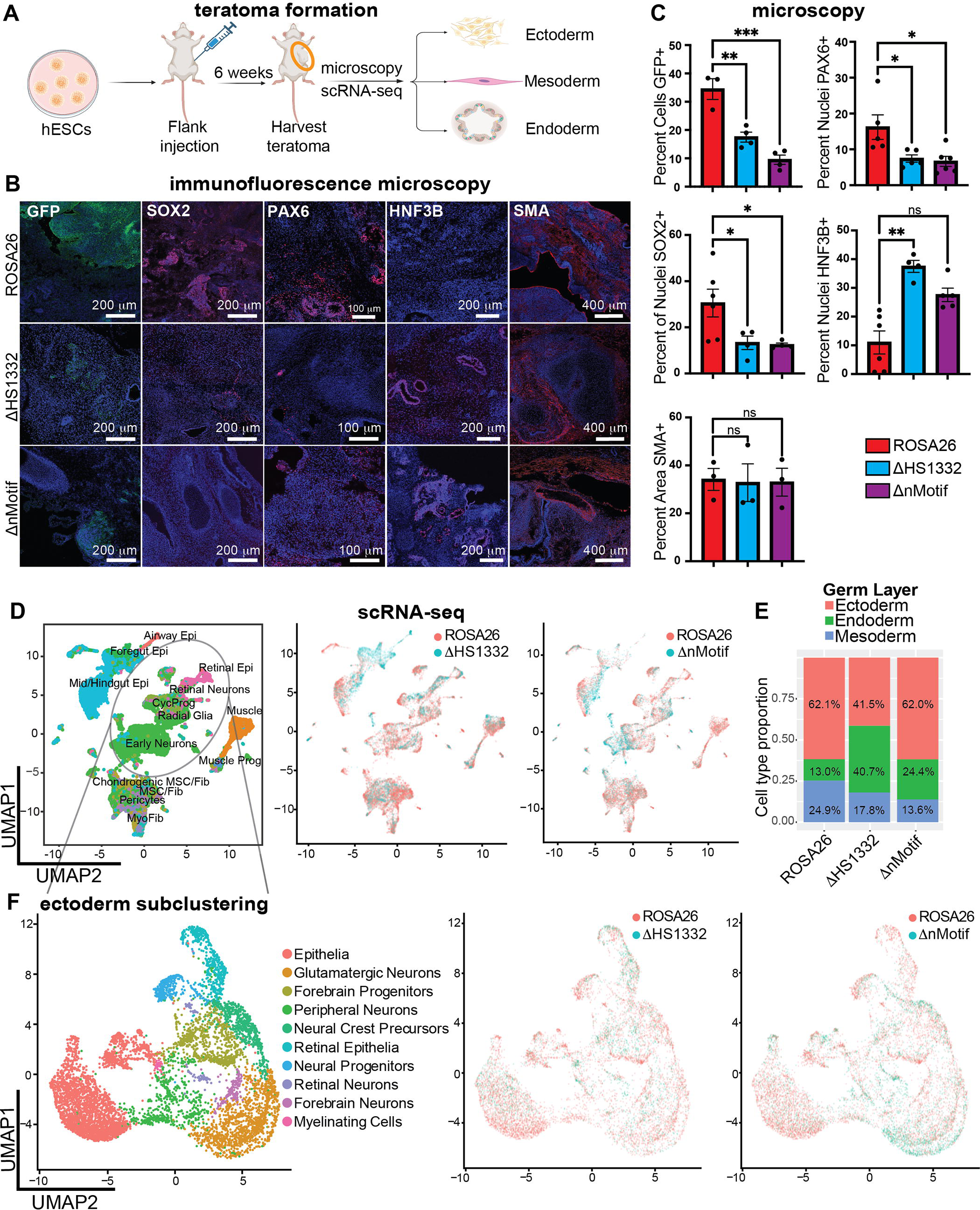
Loss of *HS1332* or CTCF-mediated 3D organization of the TAD impairs neuroectodermal development in teratomas. (A) Graphical representation of teratoma formation experiments. (B) Immunofluorescence microscopic analysis of *ROSA26*, Δ*HS1332*, and Δ*nMotif* teratomas harvested after 6 weeks. Representative images are shown for GFP, SOX2, PAX6, HNF3B, and smooth muscle actin (SMA). (C) Quantification of teratoma immunofluorescence markers: GFP (n=4 biological replicates, one-way ANOVA F_2,8_=28.84, p=0.0002; Tukey’s *post hoc* multiple comparison test: *ROSA26* vs Δ*HS1332* **p < 0.01, *ROSA26* vs Δ*nMotif* ***p < 0.001); PAX6 (n=5 biological replicates, ANOVA F_2,13_= 6.030, p=0.0140, Tukey’s *post hoc* multiple comparison test: *ROSA26* vs Δ*HS1332* *p<0.05, *ROSA26* vs Δ*nMotif* *p<0.05); SOX2 (n=4 biological replicates, ANOVA F_2,11_=4.934, p=0.0295; Tukey’s *post hoc* multiple comparison test: *ROSA26* vs Δ*HS1332* *p < 0.05, *ROSA26* vs Δ*nMotif* *p<0.05); HNF3B (n=5 biological replicates, ANOVA F_2,12_=5.297, p=0.0100; Tukey’s *post hoc* multiple comparison test: *ROSA26* vs Δ*HS1332* **p<0.01, *ROSA26* vs Δ*nMotif* p<0.05); SMA (n=3 biological replicates, F_2,6_=0.09587, p=0.9099; Tukey’s *post hoc* multiple comparison test: *ROSA26* vs Δ*HS1332* p>0.05, *ROSA26* vs Δ*nMotif* p>0.05). ns, not significant. (D) Single-cell RNA sequencing data from teratomas in UMAP representations. Clusters are labeled by cell identity (left). A comparison of *ROSA26* vs Δ*HS1332* (center) and *ROSA26* vs Δ*nMotif* (right) clusters with color coding is shown. Epi, epithelium; CycProg, cycling progenitors; Prog, progenitors; MSC, mesenchymal stem cells; Fib, fibroblasts; Myofib, myofibroblasts. (E) Bar graph representing the proportion of cells sequenced in each condition falling into an ectodermal, endodermal, or mesodermal identity. (F) scRNA-seq UMAP representation in which only ectodermal clusters from the prior UMAP (D) were extracted and re-clustered with identities assigned (left). Comparisons of ectodermal sub- clustering in *ROSA26* vs Δ*HS1332* (center) and *ROSA26* vs Δ*nMotif* (right) with color coding are shown.

To assess cell fates within teratomas from the *ROSA26*, Δ*HS1332* and Δ*nMotif* conditions in more detail, we carried out single-cell (single nucleus) RNA-seq (scRNA-seq) analysis. The analysis included 11958 cells in ROSA26, 5636 cells in Δ*HS1332*, and 6108 cells in Δ*nMotif* groups passing quality controls from at least three different teratomas in each experimental group (Figure 3D). Utilizing a prior teratoma scRNA-seq dataset encompassing 179,632 cells^52^ as a reference atlas, we identified 15 independent cell clusters encompassing all three germ layers. The teratomas generated from Δ*HS1332* hESC cells contained 41.5% ectodermal cells compared to 62.1% and 62.0% in the *ROSA26* and Δ*nMotif* conditions respectively (Figure 3E). These findings suggested that deleting the enhancer impaired neuroectodermal development with a concomitant increase in endodermal lineages.

### Impaired forebrain specification

While the Δ*HS1332* condition produced a lower proportion of ectodermal tissues as evidenced by immunofluorescence (Figure 3B), H&E staining (Figure S6A,B), and by scRNA-seq (Figure 3E), there remained a substantial neuroectodermal population. Within this neuroectodermal population, analysis of scRNA-seq data showed that the reduction in *SOX2* transcript in the Δ*HS1332* and Δ*nMotif* conditions was accompanied by decreases in *NOTCH1* mRNA, a transcriptional target of *SOX2*, as well as in *HES* and *HEY* genes that represent transcriptional targets of Notch signaling (Figure S7A,B). To identify differences within neuroectodermal lineages, we extracted the cells within the ectodermal populations of each condition from the scRNA-seq dataset, and subsequently reclustered them into ten distinct clusters comprising multiple neuronal and glial cell types (Figure 3F). We then compared the Δ*HS1332 and* Δ*nMotif* neuroectoderm to that of the *ROSA26* condition with Gene Ontology (GO) analysis using ShinyGO (Figure 4A). The major GO terms that dropped out in the Δ*HS1332* condition related to axonal development, neurogenesis and forebrain development, while in the Δ*nMotif* condition the majority of depleted GO terms related to synaptic organization and axon development^53^ (Figure 4A). In line with the depletion of GO terms relating to forebrain development, the forebrain transcription factors *FOXG1*^54,55^ and *EMX2*^56,57^ were depleted in forebrain progenitors from Δ*HS1332* and Δ*nMotif* teratomas (Figure 4B). Expression of these two transcription factors appeared dysregulated in forebrain neurons from the Δ*HS1332* and Δ*nMotif* conditions (Figure 4B), suggesting defects in neuronal maturation. These findings suggest that the *HS1332* enhancer and the associated chromatin loop contribute to neuralization, and particularly forebrain development.

**Figure 4.**
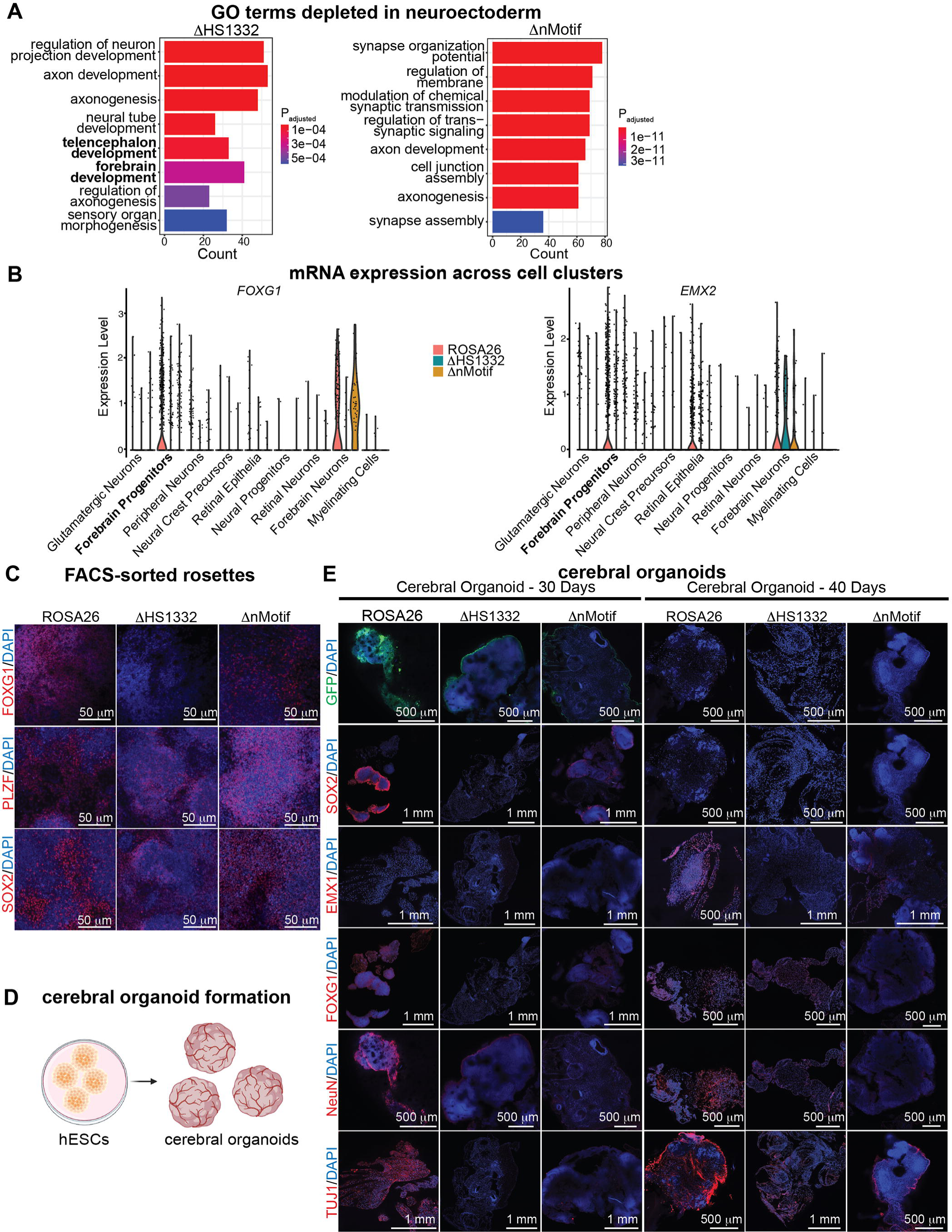
Depletion of forebrain identity by excision of *HS1332*. (A) Gene Ontology (GO) terms depleted in Δ*HS1332* and ΔnMotif vs ROSA26 teratomas in scRNA- seq analysis. (B) Violin plots of expression of *FOXG1* (left) and *EMX2* (right) transcripts across ectodermal clusters in scRNA-seq datasets. (C) Immunofluorescence microscopy of FACS-sorted GFP-positive rosettes stained for FOXG1, PLZF, and SOX2 in hESC derived rosettes. PLZF is a TF marker of rosette progenitors.^50,76^ (D) Graphical representation of cerebral organoid generation from hESCs. (E) Organoids from the *ROSA26*, Δ*HS1332*, and Δ*nMotif* conditions at the 30 (left) and 40 (right)-day time-points were stained for the forebrain markers FOXG1 and EMX1, neuronal markers NeuN and TUJ1, SOX2 and GFP.

To further investigate the forebrain phenotype observed in teratoma formation assays, hESCs were differentiated to rosettes using dual SMAD inhibition.^58^ Immunofluorescent staining and imaging of Δ*HS1332* rosettes demonstrated a near complete lack of FOXG1 immunostaining (Figure 4C), confirming the findings in teratomas.

To collect additional evidence for the forebrain phenotype, we tested the function of the enhancer in cerebral organoids^59–62^ from *ROSA26*, Δ*HS1332* and Δ*nMotif* hESCs (Figure 4D). Organoids were analyzed at the 30 and 40-day time-points. Immunostaining for SOX2 and GFP, from the *Hes5::GFP* reporter, were significantly reduced in the Δ*HS1332* and Δ*nMotif* conditions at 30 days, but present in the *ROSA26* condition. TUJ1 and RBFOX3 (NeuN)-positive neuronal fates were significantly reduced in the Δ*HS1332* and Δ*nMotif* conditions at both time-points (Figure 4E). Furthermore, immunostaining for forebrain transcription factors FOXG1 and EMX1 showed depletion in ΔHS1332 organoids, and in Δ*nMotif* organoids to a lesser extent. These observations support the hypothesis that the distant *SOX2* enhancer, and the associated 3D chromatin loop to a lesser extent, are particularly crucial to telencephalic development.

### HS1332 enhancer activation does not depend on chromatin conformation changes

The fact that activation of the enhancer and formation of the chromatin loop within the *SOX2* TAD are developmentally co-regulated and specific to neuroectodermal fates raises the possibility of causality linking the two processes. To answer this question, we performed ChIP-seq for the activating chromatin modification H3K27ac and the facultative repressive modification H3K27me3, as well as 4C using a bait in the *SOX2* promoter, in hESCs and rosettes. In hESCs from all conditions, the enhancer element was neither activated nor repressed (Figure 5A,B), but in the Δ*HS1332* condition the *SOX2* TAD had overall diminished H3K27ac deposition and a locus centromeric to *HS1332* was heavily decorated with the H3K27me3 modification (peak 1 in Figure 5B and Figure S8). At the rosette stage, both the control *ROSA26* condition and the Δ*HS1332* and Δ*nMotif* conditions showed decoration of the enhancer element with the activating H3K27ac modification and lack of the repressive H3K27me3 mark (Figure 5A,B). Interestingly, the *SOX2* TAD harbored increased H3K27me3 in the rosettes of the Δ*HS1332* condition at two loci telomeric to the enhancer (peaks 2 and 3 in Figure 5B and Figure S8). Peak 2 in Figure 5B localizes in the immediate vicinity of the long non-coding RNA *LINC01994*.^35^ 4C-seq detected little long-range contact between the enhancer region and the *SOX2* promoter in all hESCs. In contrast, the robust long-range contact seen in *ROSA26* rosettes was significantly diminished in the Δ*nMotif* condition (Figure 5C,D). Collectively, these data suggest that enhancer activation is not coupled to establishment of the chromatin loop, since it is observed in both the *ROSA26* and Δ*nMotif* conditions, and that the enhancer regulates activating and repressive epigenetic modifications in the *SOX2* TAD.

**Figure 5.**
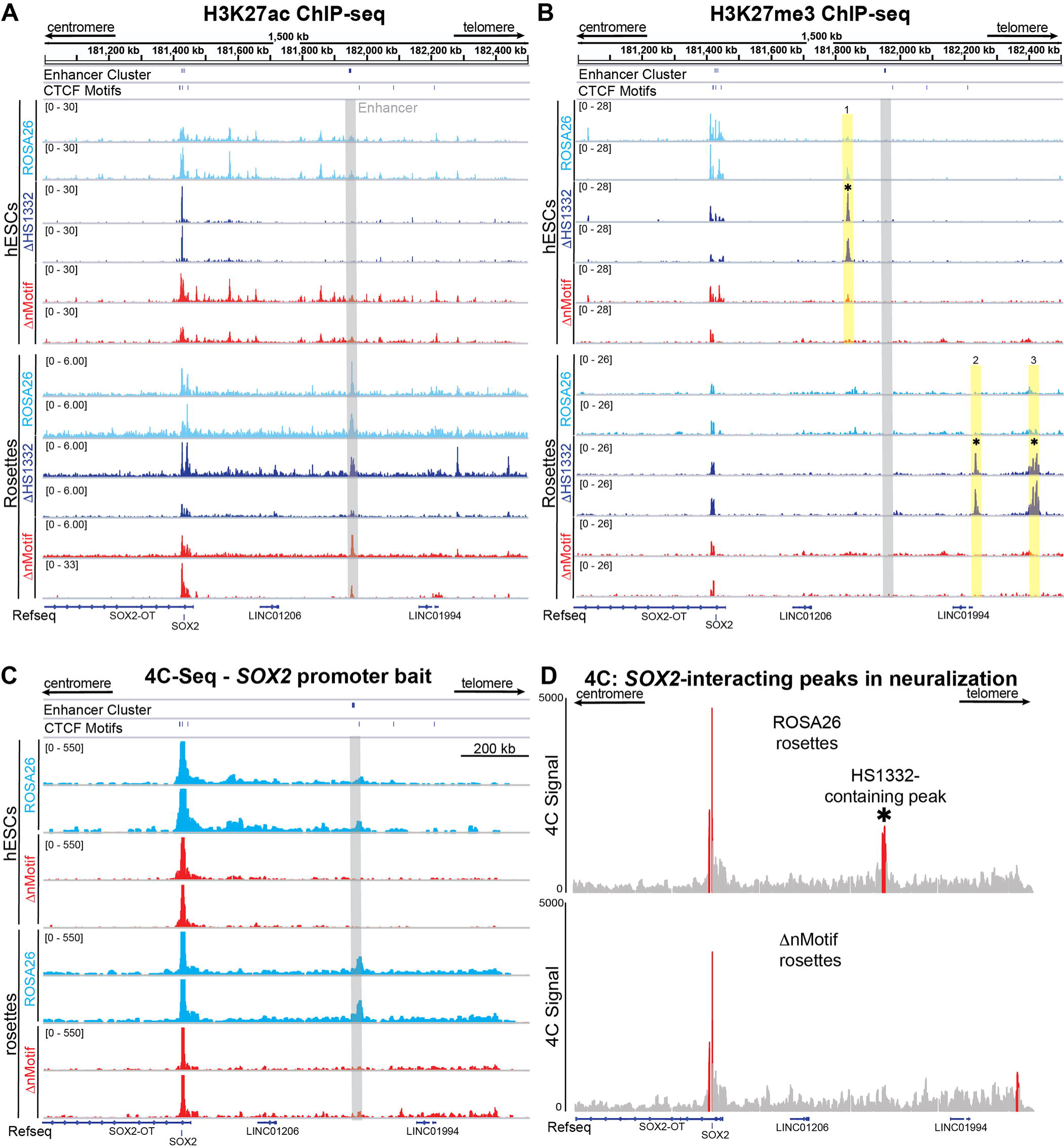
Effects of *HS1332* excision and 3D chromatin loop perturbation on chromatin modifications and 3D architecture in the *SOX2* TAD. (A) Genome browser view of H3K27ac ChIP-sequencing in the *SOX2* TAD in hESCs using RPGC-normalized signal track and rosettes using input-normalized ppois signal track. The gray highlight identifies the location of the *HS1332* enhancer. (B) Genome browser view of RPGC-normalized H3K27me3 ChIP-sequencing in the *SOX2* TAD in hESCs and rosettes. Additional data for peaks 1-3 are shown in Figure S8. The gray highlight identifies the location of the *HS1332* enhancer, while the yellow highlights show areas of increased H3K27me3 deposition in the Δ*HS1332* condition. (C) 4C-sequencing with a *SOX2* promoter bait in hESCs and rosettes. The gray highlight identifies the location of the *HS1332* enhancer. (D) Significant interacting peaks, as determined by Benjamini-Hochberg multiple testing procedure (red), in 4C-sequencing data of rosettes from the *ROSA26* and Δ*nMotif* conditions (*p<0.05).

### Disruption of the chromatin loop after NSC specification replicates SOX2 expression changes and differentiation

The experiments above tested effects of perturbing enhancer activation and chromatin looping during specification of neuroectoderm. We next tested the importance of this 3D chromatin organization in already specified human NSCs. We established cultures of hESC-derived NSCs in epidermal growth factor (EGF) and fibroblast growth factor 2 (FGF2)-containing medium and then excised the CTCF *nMotif* using a dual gRNA CRISPR-Cas9 approach. A NSC line expressing a non-targeting (NT) gRNA was used as control. Excision of the targeted motif (90 bp) was verified by PCR amplification and Sanger sequencing of the resulting product (Figure 6A). Excision of the motif decreased *SOX2* transcript levels and impacted the growth and viability of NSC cultures (Figure 6B-D). Most importantly, abrogation of this chromatin loop prevented NSC differentiation to neurons and facilitated differentiation to astrocytes (Figure 6E,F). These findings suggested a critical function for 3D chromatin organization in this TAD in *SOX2* transcription, NSC multipotency and neuroglial differentiation.

**Figure 6.**
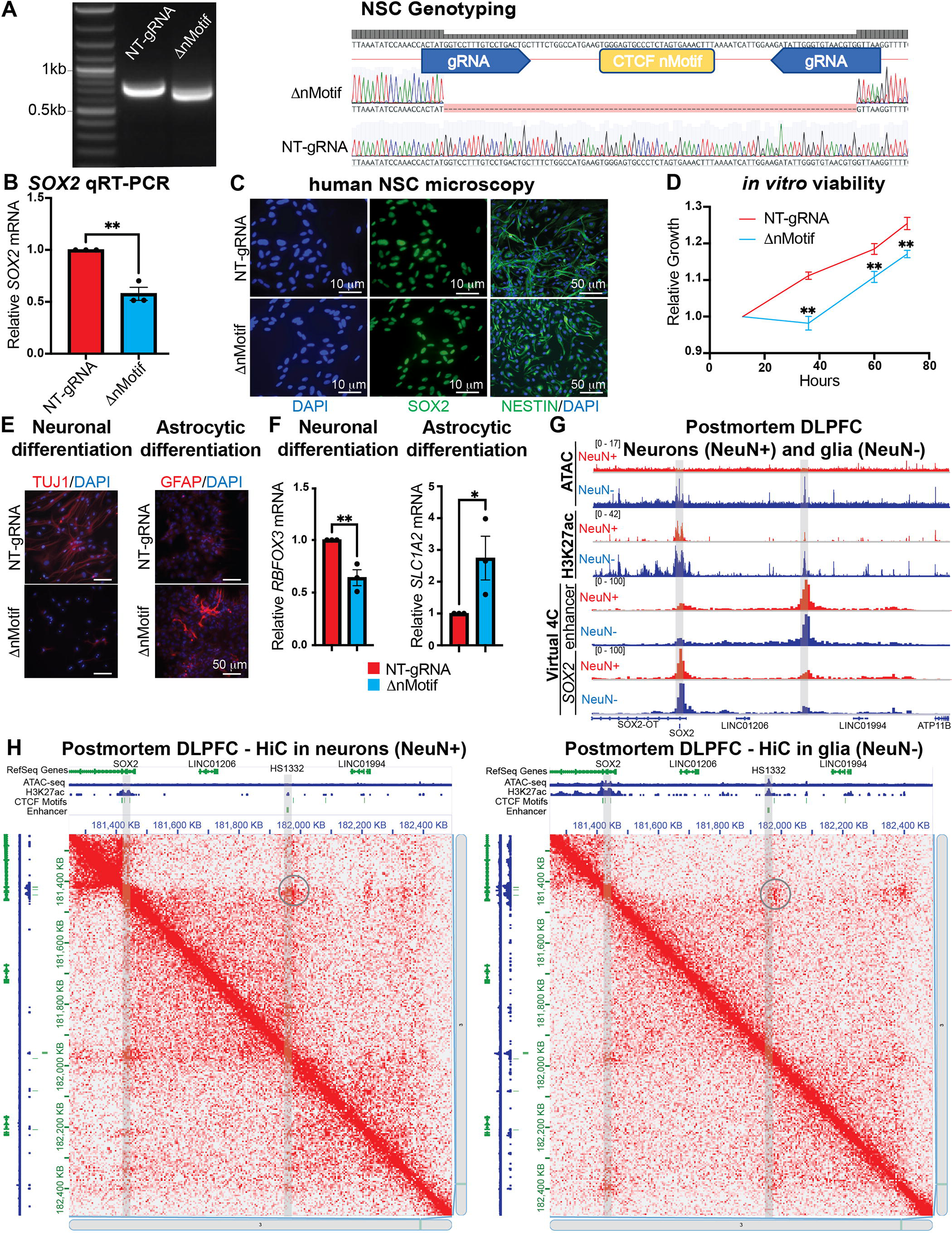
Loss of *SOX2* local chromatin organization affects *SOX2* expression and neuroglial differentiation in mature NSCs. (A) Agarose gel electrophoresis of PCR product containing the n*Motif* domain in control (NT-gRNA) vs Δ*nMotif* NSCs (left). The Sanger sequencing results of the two PCR products are also shown (right). (B) qRT-PCR for *SOX2* transcript in control and Δ*nMotif* NSCs (n=3 biological replicates, t-test **p<0.01) (C) Immunofluorescence microscopy for SOX2 and NESTIN in control vs Δ*nMotif* NSCs. (D) Growth/viability measurement by WST8 absorbance assay in control and Δ*nMotif* NSCs (n=3 biological replicates, two-way ANOVA n=3, F_3,6_=32.1 p<0.01; Sidak’s multiple comparisons test **p<0.01 at indicated timepoints). (E) Immunofluorescence microscopy for TUJ1 and GFAP in neuronal and astrocytic differentiation of control and ΔnMotif NSCs. (F) qRT-PCR of neuronal transcript *RBFOX3* and astrocytic transcript *SLC1A2* after neuronal and astrocytic differentiation of control and Δ*nMotif* NSCs (n=3 biological replicates, t-test p=0.0091 and p=0.0171, respectively). (G) Genome browser view of *SOX2* TAD comparing the following assays in NeuN+ or NeuN- cells isolated from postmortem DLPFC in PsychEncode: ATAC-Sequencing, H3K27ac ChIP- sequencing, *SOX2* promoter virtual 4C-sequencing, and *HS1332* enhancer virtual 4C- sequencing. (H) HiC plots of the *SOX2* TAD in NeuN+ or NeuN- cells of postmortem DLPFC show preservation of 3D architecture in adult neurons and glia.

The relevance of this chromatin loop and enhancer-promoter contact to neuroglial specification was corroborated by analysis of PsychEncode ATAC-seq and H3K27ac data from postmortem adult dorsolateral prefrontal cortex (DLPFC).^63^ Our analysis showed that, while both neurons (NeuN+) and glia (NeuN-) retained the chromatin loop apposing the enhancer and *SOX2* locus, the two lineages exhibited differential chromatin accessibility and euchromatic modifications (Figure 6G,H).^63,64^ Indeed, NeuN- glial lineages showed greater chromatin accessibility within the *SOX2* TAD, with both the *SOX2* gene locus and the *HS1332* enhancer bearing more H3K27ac euchromatic modifications than NeuN+ neuronal lineages, a finding that may be related to reported persistence of *SOX2* expression in classes of adult astrocytes.^65^ These findings support a model in which the chromatin loop is required for NSCs to generate neuroglial progeny, and also retained in the progeny despite differences in chromatin state between neuronal and glial cells.

## DISCUSSION

Differentiation requires dynamic control of gene expression, for which enhancer recruitment is a core determinant.^12,66,67^ The recruitment of enhancers is frequently associated with the induction of genes that are quiescent in prior development stages, thus ascribing to them important functions during developmental transitions. Understanding the activity of these regulatory mechanisms can aid in dissecting their role in development and in improving the efficiency of directed stem cell differentiation to specific lineages for research and therapeutic purposes.

SOX2, alongside OCT4 and NANOG, forms a core pluripotency TF network in ESCs that acts to sustain transcription of all three genes, while also maintaining pluripotency and suppressing differentiation programs.^10,11,13,14^ In the developing nervous system, however, where OCT4 and NANOG are absent, *SOX2* transcription is maintained via reliance on other incompletely understood mechanisms, including other TFs and enhancer elements.^68–70^ The importance and functional roles of such neuroectodermal enhancers acting on *SOX2* have only been partially explored.^33,34,38^

This study points to the importance of dynamic reconfiguration of chromatin organization during development. Within the TAD where *SOX2* resides, the chromatin landscape undergoes significant changes during the specification of neuroectoderm. We discovered that in hESCs the *SOX2* locus does not form strong 3D contacts with other elements within the TAD and the *HS1332* enhancer is silent but not repressed, as suggested by lack of facultative repressive H3K27me3 modifications – findings similar to observations in mouse models^37,43,44^. Upon differentiation to neuroectodermal stem cell fates, however, *HS1332* and the neighboring *HS1132b* loci are decorated with euchromatic activating H3K27ac modifications and a 3D chromatin loop with the *SOX2* gene is established, likely mediated by CTCF binding to motifs adjacent to the enhancer and the gene. Perturbation of the enhancer or its 3D contact with the *SOX2* gene impairs the formation of neuroectoderm, and the forebrain in particular. Similar phenotypes have been reported with perturbation of proximal *Sox2* enhancer clusters in mouse models^34^, raising the possibility of multiple regulatory elements for neural *SOX2* transcription in mammals. Mechanistically, the establishment of the 3D contact and the activation of the enhancer occur independently of each other even though they coincide spatiotemporally during neural development. This is evidenced by the finding that the enhancer still manifests activating chromatin modifications when the 3D chromatin loop is disrupted by excision of the adjacent CTCF motif.

Multiple lines of evidence suggest that this coordinated activation of the enhancer and the chromatin loop in neural lineages are crucial developmental events. First, the genomic sequences representing the enhancer are highly conserved from fish to primates. Second, the enhancer element manifests the highest amount of euchromatic modifications within the *SOX2* TAD in neuroectoderm besides the *SOX2* gene. Third, the 3D contact between the *SOX2* locus and enhancer is the strongest within the TAD and highly conserved in evolution. Finally, this 3D chromatin architecture is preserved in mature neuronal and glial lineages in the human brain, indicating either “imprinting” or a lasting functional role for it after neuroglial differentiation. Collectively, these observations suggest that this developmentally co-regulated enhancer activation and contact with the *SOX2* gene is an evolutionarily ancient mechanism essential to neuroectodermal development and neuroglial specification. This hypothesis received strong support from our perturbation experiments.

Our data suggest a role for this 3D interaction not only during developmental transitions from pluripotency to neuroectodermal fates, but also after establishment of the neural lineage. Indeed, ablation of the CTCF motif adjacent to the enhancer in NSCs prevented their differentiation to neurons, a central property of NSC multipotency. This finding, along with the observation that this 3D loop and the TAD topology are preserved in mature neurons and glia, suggests that enhancer-promoter interactions and 3D chromatin architecture do not just govern specification of stem cells, but are also critical in the maintenance of stem cell states.

Of particular interest is the finding that hemizygous deletion of the enhancer produced stronger phenotypes than homozygous deletion of the CTCF motif responsible for the 3D contact between the *SOX2* gene and the enhancer. Although our 4C sequencing showed that deletion of the particular CTCF motif substantially reduced the strength of the related chromatin loop, it is possible that the residual infrequent 3D contact suffices for the enhancer to elicit effects on *SOX2* transcriptional regulation via a “kiss-and-run” mechanism^71^. An alternative explanation is that this enhancer regulates multiple elements within the TAD beyond just the *SOX2* gene. Consistent with this theory is the fact that deletion of the enhancer results in repression of other non-coding sequences within the *SOX2* TAD (peaks 2 and 3 in Figure 5B), including long non-coding RNAs.^35^

The reason that we were able to achieve only a hemizygous but not a homozygous deletion of the *HS1332* is unclear. The size of the genomic sequence targeted for deletion, 2.7 kb, may be a contributing factor in reducing efficacy of the excision. Alternatively, it is possible that homozygous deletion of the enhancer is deleterious to hESCs. Aligned with this hypothesis is the observation that, while our assays indicated no effects of the hemizygous *HS1332* deletion on hESC self-renewal and pluripotency markers, our ChIP-sequencing revealed increased H3K27me3 deposition in a non-coding sequence in the *SOX2* TAD (peak 1 in Figure 5B) at the hESC stage. Collectively, these observations raise the possibility that the *HS1332* enhancer may exert effects on several sequences within the *SOX2* TAD besides the *SOX2* gene itself in both neuroectodermal lineages and potentially hESCs as well.

The essentiality of SOX2 to the development of the central nervous system, and neurons in particular, has been documented both experimentally and with human genetic studies. In our hands, SOX2 haploinsufficiency arising from *HS1332* deletion or excision of the adjacent CTCF motif resulted in impaired specification of neural progenitors and a profound decrease in neuronal differentiation. Our findings are in line with previous mouse models showing that neural progenitors with *SOX2* hypomorphic genetic alterations produce neurons less efficiently,^19,20^ and with clinical data from patients with *SOX2* haploinsufficiency leading to anophthalmia or microphthalmia, brain malformations and learning disabilities, as well as esophageal, genital and endocrinologic developmental abnormalities.^72–74^ Similarly, haploinsufficiency of *FOXG1*, a TF critical to forebrain development^55^ whose expression is significantly reduced in our hands after deletion of the *HS1332* enhancer or perturbation of the associated chromatin loop, is associated with brain malformation, microcephaly, mental retardation and Rett syndrome.^72,75^

Our work has important implications for human stem cell and regenerative biology. Understanding the complex genomic mechanisms that govern developmental transitions in humans is crucial to improving protocols for directed differentiation from existing stem cell platforms. As an example of the translational utility of our findings, one could test whether direct activation of the enhancer element facilitates the generation of human neurons with a forebrain identity. Of equal interest and translational potential is the fact that SOX2 is expressed in tumors deriving from tissues whose development depends on SOX2, such as glioma, lung, and esophageal adenocarcinoma.^28–30,35^ Examining how enhancers such as the one we investigated function in the context of cancer may shed light on the oncologic regulation of *SOX2* expression and offer targets for therapy.

## RESOURCE AVAILABILITY

### Lead contact

Further information and requests for resources and reagents should be directed to and will be fulfilled by the lead contact, Dimitris G. Placantonakis.

### Materials availability

Cell lines are available upon request.

### Data and code availability

• Single-cell RNA-seq, ATAC-seq, HiC, and 4C-seq data have been deposited at GEO (GSE318965, GSE318999, GSE320454) and are publicly available.

• Any additional information required to reanalyze the data reported in this paper is available from the lead contact upon request.

## ACKNOWLEDGMENTS

This work was supported by NIH R01NS124920 (DGP), NYSTEM (NY State Stem Cell Science) IIRP C32595GG (DGP), NIH R01NS102665 (DGP), NIH R21NS126806 (DGP), NIH F30CA247418 (DB), NIH R35GM122515 (JS), NIH P01CA229086 (JS), NIH R01AG075272 (TL), NIH R01CA260028 (TL), and NIH P30CA016087 (NYU Perlmutter Cancer Center Support Grant). We acknowledge the PsychEncode Consortium for sharing data from human brain specimens.

## AUTHOR CONTRIBUTIONS

Conceptualization, DB and DGP; methodology, DB, SW, JS, DGP; investigation, all authors; writing— original draft, DB and DGP; writing—review & editing, all authors; funding acquisition, DB, DGP; resources, MS, TL, AS, JS, DGP; supervision, DGP.

## DECLARATION OF GENERATIVE AI AND AI-ASSISTED TECHNOLOGIES

While preparing this work, the author(s) did not utilize any generative AI or AI-assisted technologies.

## METHODS

### H9 hESC culture

H9 (WA09) human embryonic stem cells (hESCs) stably nucleofected with a *Hes5::GFP* reporter were cultured as colonies on a layer of mouse embryonic fibroblasts (MEFs, Invitrogen) in hESC medium, consisting of 1:1 DMEM/F12 (Invitrogen), 20% KSR (Invitrogen), 5 ng/mL FGF2 (R&D), 2 mM glutamine (Invitrogen), 0.1 mM non-essential amino acids (Invitrogen), and 0.1 mM β-mercaptoethanol (Invitrogen).^47^ Cultures were routinely groomed of differentiating colonies and fed daily. Passaging was performed with dispase (Invitrogen) or by mechanically picking colonies. Dissociation to single cells was performed with Accutase (Thermo Fisher) and subsequent culture of cells in hESC medium supplemented with 10 μM Y-27632 ROCK inhibitor (MedChemExpress) for the first 24 hours. All hESC experimentation was approved by ESCRO (protocol #14-00267) at NYU Grossman School of Medicine. Cells were cultured in humidified cell culture incubators at 37 °C with room air balanced with 5% CO_2_.

### HEK293T cell culture and lentivirus production

Human embryonic kidney 293 T cells (HEK293T, Takara, Cat# 632180) were cultured in Dulbecco’s Modified Eagle’s Medium (DMEM, Gibco) supplemented with sodium pyruvate (Gibco) and 10% fetal bovine serum (FBS; Peak Serum) at 37 °C and 5% CO_2_ in humidified room air. HEK293T cells were co-transfected with plasmid DNAs consisting of the lentiviral transfer vector, as well as the pMD2.G and psPAX2 packaging plasmids using Lipofectamine 2000 (Invitrogen). Medium was harvested for 3 consecutive days with Lenti-X Concentrator (Takara) added at 1:4 and stored at 4 °C. Virus-containing medium was centrifuged at 1500xg for 45 minutes at 4 °C to concentrate the viral particles. Viral particles were resuspended in Opti-MEM (Thermo Fisher), aliquoted, and stored at -80 °C until use.

### CRISPR genome editing

Flanking gRNAs for target motifs or enhancers were designed using Benchling with on-target and off-target scores calculated according to published methods. Dual gRNA inserts were cloned into LentiCRISPRv2-mCherry in accordance with previously published methods.^77,78^ In short, oligos containing both gRNAs (Table 1) were acquired from Eurofins Genomics and then PCR-amplified to create a fragment which was subcloned into a *U6* promoter-containing donor construct. The resulting dual promoter and dual gRNA fragment was cloned into one of the previously listed LentiCRISPRv2 constructs. Successful cloning was confirmed by diagnostic digest with XhoI and NotI (New England Biolabs), followed by Sanger sequencing of the gRNA insert. Lentivirus was generated in HEK293T cells by cotransfection with the packaging plasmids pMD2.G and psPAX2 as previously described.^79^ hESCs or NSCs were transduced with lentivirus at a multiplicity of infection of 1. mCherry-positive hESCs were FACS-sorted in hESC medium supplemented with 10 μM Y-27632 and expanded in MEF cultures. To obtain hESC clones with the desired CRISPR modifications, we dissociated hESC colonies to single cells using Accutase (Thermo Fisher) and the resulting suspension was filtered through a 40 μm filter to ensure a single cell suspension. Individual cells were seeded on MEFs in 24 well tissue culture plates at 1,2,4 and 5 cells per well. Only the 5 cell/well condition produced colonies at a rate of 1 for every 24 wells seeded. mCherry-positive NSCs were FACS-isolated and pooled cell cultures were grown for two weeks before genotyping.

**Table 1.**
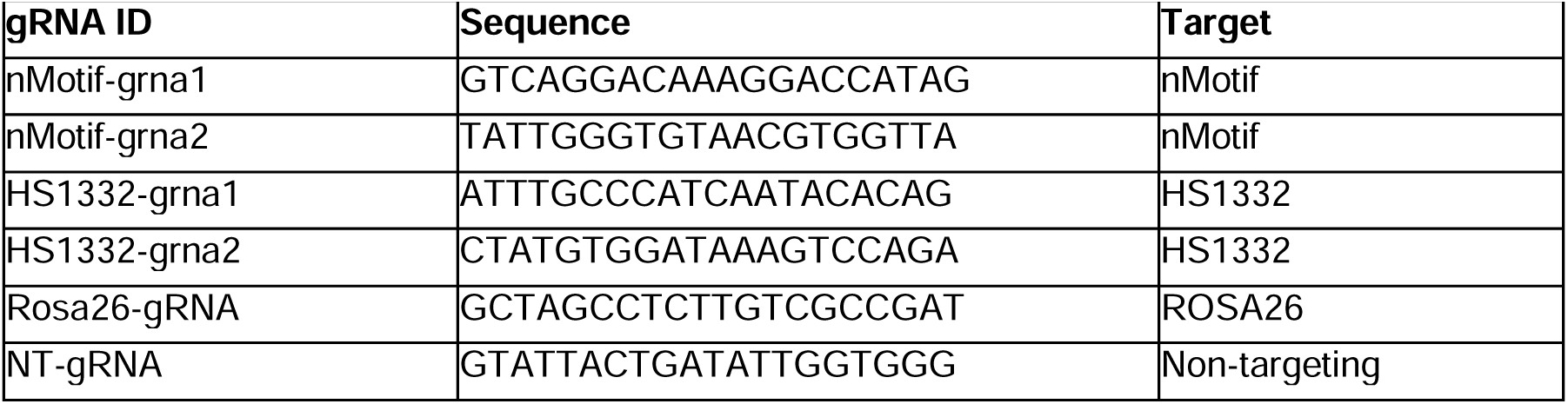
gRNAs used for targeted CRISPR deletions.

### Genotyping of modified cell lines

Human ESCs and NSCs selected for expression of lentiCRISPR dual gRNA constructs were genotyped using primers flanking the modified loci (Table 2). Resulting PCR products were run on 1% agarose gels containing SYBR Safe (Thermo) for visualization and imaged using an Invitrogen iBright. PCR products in modified cell lines were extracted from agarose gels and Sanger-sequenced to verify deletion at the predicted motifs. Human ESC clones and NSCs with verified deletions were then used for all subsequent analyses.

**Table 2.**
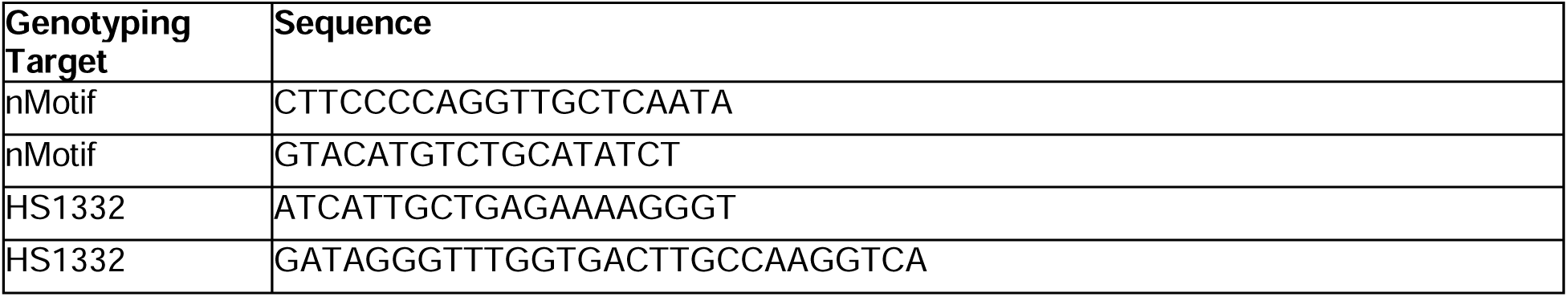
Primers for Genotyping Deletions.

### Directed hESC to rosette-NSC differentiation *in vitro*

Human ESC to rosette-NSC (rosette) differentiation was performed, as previously described.^58^ Briefly, hESCs were plated on Geltrex-coated plates at ∼50,000 cells/cm^2^ and grown under feeder-free conditions in Pluristem Human ES Cell Medium (Millipore) until reaching ∼80% confluence. Medium was then changed first to KSR medium [Knockout-DMEM, 15% knockout serum replacement (KSR), 2 mM L-glutamine, 1X nonessential amino acids (NEAA), and 0.1% β-mercaptoethanol] supplemented with 10 μM SB-431542 (stemMACS) and 100 nM LDN-193189 (Fisher) to achieve dual SMAD inhibition. KSR concentration was gradually reduced over 10 days in exchange for N2 medium: on day 2, medium was changed to 75% KSR medium + 25% N2 Medium [1:1 DMEM/F12 (Invitrogen), N2 supplement 1X, and 1.6 g/L glucose], again supplemented with SB-431542 and LDN-193189; on days 4, 6, and 8, medium was changed with a 25% decrease in KSR medium composition every two days, until on day 8 the medium consisted of 100% N2 with SB-431542 and LDN-193189.

### NSC culture and differentiation

Human ESC-derived NSCs were cultured in DMEM/F12 medium (Thermo) supplemented with N2 supplement 1X (Fisher), B27 supplement 0.1X (Fisher), insulin (Sigma), glucose (1.6 g/L) (Fisher), 20 ng/mL EGF (R&D Systems) and 20 ng/mL FGF2 (R&D Systems). Cells were cultured on plates coated for 24 hours with 0.025% poly-L-ornithine (Gibco) followed by 2 hours of coating with 10 μg/mL of laminin in PBS (Gibco). Medium was changed every 48 hours. Cells were passaged upon reaching confluence with Trypsin (Invitrogen) and Defined Trypsin Inhibitor (Invitrogen) onto freshly coated plates at a 1:2 passaging dilution. Low-passage cell cultures were used for all experiments.

### Neuroglial differentiation

For neuronal differentiation, rosettes or NSCs maintained in EGF/FGF2 were cultured in N2 medium with 20 ng/mL BDNF (R&D) and 200 nM Ascorbic Acid (Sigma). Medium and growth factors were replaced every two days for 30 days before analysis. Astrocytic differentiation was performed with 4% FBS (Invitrogen) in N2, with fresh medium changed every two days for 42 days. Cells were cultured in humidified cell culture incubators at 37 °C balanced with 5% CO_2_. Biological replicates represent differentiations performed on different days using newly seeded rosettes or NSCs maintained in EGF/FGF2.

### Serum-free embryoid body (SFEB) culture

Mature hESC colonies grown on MEF plates were dissociated using dispase (Sigma). The resulting colonies were collected, resuspended in APEL2 serum-free media (Stemcell Technologies), and triturated by pipetting 3 times to break up mature colonies into smaller colonies. Around 15-20 colonies were plated on low-attachment 24-well plates (Thermo Fisher) in 500 ml of APEL2 medium.^49^ Medium was refreshed every 5 days with new APEL2 medium and SFEBs were monitored by wide-field microscopy to assess GFP fluorescence.

### hESC teratoma formation assay

Male NSG mice at 6 to 8 weeks of age were anesthetized using an anesthetic of ketamine and xylazine diluted in PBS at 100 mg/kg and 20 mg/kg, respectively. After induction of anesthesia, each flank was sterilized with 70% ethanol and shaved. Subsequently, 1x10^6^ dissociated hESCs in 50 mL of a 1:1 mixture of DMEM/F12 (Invitrogen) and Matrigel (Corning) was drawn into a 17-gauge needle and injected into the flank. Teratomas were allowed to grow for 6 weeks. Teratoma tissue was harvested following euthanasia through injection with 50 mg/kg of pentobarbitol (Euthasol, Virbac). Procedures were performed according to IACUC protocol number 160403 at NYU Grossman School of Medicine.

### Cerebral organoid culture

Cerebral organoids were made using the forebrain cerebral organoid kit from StemCell Technologies. To generate the initial embryoid bodies, low-passage hESC colonies were dissociated to single cells using Gentle Dissociation Reagent and resuspended in Basal Medium 1 with Supplement A containing 10 μM Y-27632 at a concentration of 150,000 cells/mL. One hundred μL of the cell suspension was plated in ultra-low attachment round-bottom 96-well plates (Corning) and incubated for five days, with fresh medium added every other day. At the 5-day mark, embryoid bodies were transferred to low-attachment 24-well plates and grown for 2 days in induction media composed of Basal Medium 1 and supplement B. At day 7, embryoid bodies were embedded in Matrigel on a sterile embedding surface, transferred into low-attachment 6-well plates, and maintained in expansion media for 3 days. Beginning on day 10 and continuing through day 30 or 40, cerebral organoids were maintained in maturation media, which was changed every 3 days until organoids were harvested.

### Chromatin immunoprecipitation (ChIP) sequencing

ChIP-Sequencing libraries were prepared according to the ChIPmentation protocol previously published.^80^ In short, hESCs or their rosette progeny were fixed in 1% paraformaldehyde (PFA) for 10 minutes at room temperature (RT) and quenched with a final concentration of 0.125 M glycine for 5 minutes. Fixed cells were lysed and sonicated using a Diagenode Bioruptor for 15 cycles at 30 seconds on, 30 seconds off on the “high” sonication setting. Proper chromatin shearing with the desired fragment length in the 200-600 bp range was confirmed by agarose gel electrophoresis. Chromatin was incubated at 4 °C overnight with antibody-conjugated dynabeads. Beads were washed and chromatin was tagmented using Tn5 transposase (Illumina). DNA was de-crosslinked by proteinase K digestion and was purified using the Qiagen MinElute kit. Tagmented, de-crosslinked DNA was PCR-amplified using single barcoded primers and was sequenced with a target sequencing depth of >30 M reads for H3K27ac or CTCF ChIP, and >50M reads for H3K27me3 ChIP samples.

### Assay for Transposase-Accessible Chromatin (ATAC) sequencing

We used an improved ATAC-seq protocol (Omni-ATAC) adopted from Corces et al.,^81^ with small adaptations. After counting, 100,000 cells were collected, resuspended in 1X cold DPBS (Gibco) and centrifuged at 500xg for 5 min at 4 °C. After centrifugation, the supernatant was removed and the cell pellet resuspended in 500 μL of cold ATAC-seq resuspension buffer 1 (10 mM Tris-HCl pH 7.4, 10 mM NaCl, 3 mM MgCl_2_, 0.1% NP-40 and 0.1% Tween-20 in H_2_0). Fifty μL (50,000 cells) were transferred to a new 1.5 ml Eppendorf tube and 0.5 μL 1% Digitonin (Promega) was added by pipetting well a few times. The cell lysis reaction was incubated on ice for 3 min. After lysis, 1 mL of cold ATAC-seq resuspension buffer 2 (10 mM Tris-HCl pH 7.4, 10 mM NaCl, 3 mM MgCl_2_ and 0.1% Tween-20 in H_2_0) was added. The tube was inverted three times to mix the contents and nuclei were pelleted by centrifugation for 10 min at 500xg at 4 °C. The supernatant was carefully removed and nuclei were resuspended in 45 μL of transposition mix [25 μL 2X TD buffer (Illumina), 16.5 μL 1X PBS, 0.5 μL 1% digitonin, 0.5 μL 10% Tween-20 and 2.5 μL H_2_O) by pipetting up and down a few times. Five μL of transposase enzyme (Illumina) was then added and the transposition reaction was incubated at 37 °C for 30 min in a thermomixer with shaking at 1000 rpm. The reaction was cleaned up with MinElute Reaction Cleanup Kit (Qiagen). Transposed DNA fragments were then amplified as described previously in Buenrostro et al.^82^ Final libraries were purified with Ampure XP Beads (Beckman), checked by Tapestation High Sensitivity (Agilent) and quantified by Qubit (Life Technologies). Libraries were then sequenced with the Novaseq6000 Illumina Sequencing System with 150 bp paired-end reads.

### RNA sequencing

RNA was extracted from 5x10^6^ freshly harvested hESCs or NSCs using the Monarch Spin RNA Isolation Mini Kit (NEB). Quality was verified by an Agilent Bioanalyzer 2100. RNA-seq libraries were generated by the NYU Genome Technology Center using the Illumina Ribo-Zero Plus rRNA Depletion Kit and sequenced on the Illumina NovaSeq X with 50 bp paired-end reads. RNA-seq fastq files were processed in route “rna-star” using the Seq-n-Slide (sns) pipeline: https://igordot.github.io/sns/. Alignment of reads was performed using the hg19 assembly of the human genome. BAM alignment files were processed to BigWig using deeptools and visualized using the UCSC genome browser.^83,84^

### HiC library preparation

Human ESCs or NSCs were dissociated and counted, and 1x10^6^ cells were crosslinked with 2% methanol-free formaldehyde (Tousimis) in PBS for 10 minutes at RT. The reaction was quenched with glycine to a final concentration of 0.125 M. The cell suspension was then centrifuged and the supernatant was removed. Cells were washed twice with PBS and processed using the HiC+ kit from Arima Genomics. In short, crosslinked cells were incubated with multiple restriction enzymes to digest chromatin. Arima-HiC sequencing libraries were prepared by first shearing purified proximally ligated DNA and then size-selecting 200-600 bp DNA fragments using SPRI (Solid Phase Reversible Immobilization) beads. The size-selected fragments were then enriched using Enrichment Beads (provided in Arima HiC kit), and then converted into Illumina-compatible sequencing libraries with TruSeq adapters using the KAPA HyperPrep kit (Roche). The purified, PCR-amplified libraries underwent standard quality control using KAPA qPCR and Tapestation (Agilent) and were sequenced on the NovaSeq6000 Illumina Sequencing System with 150 bp paired-end reads.

### 4C library preparation

Cells were processed for 4C-Seq as previously described.^39^ Cells were dissociated and counted, and 1x10^7^ cells were crosslinked with 2% methanol-free formaldehyde (Tousimis) in PBS for 10 minutes at RT. The reaction was quenched with glycine to a final concentration of 0.125 M. After centrifugation, the cell pellet was resuspended in 1 mL of lysis buffer (50 mM Tris, 150 mM NaCl, 5 mM EDTA, 0.5% NP-40, 1% Triton X-100 and 1X Roche complete protease inhibitor). Cells were incubated on ice for 15 minutes and then centrifuged at 2500 RPM at 4 °C for 5 minutes. Pellets were resuspended in 360 μL of molecular grade H_2_O, 60 μL of DpnII restriction buffer and 15 μL of 10% SDS. Samples were incubated at 37 °C for 1 hour shaking at 900 RPM. One hundred fifty μL of 10% Triton X-100 was added and samples were further incubated at 37 °C for 1 hour at 900 RPM. A 5 μL sample was taken and stored for comparison with the prepared template. Nuclei were incubated overnight with 200 units of DpnII (New England Biolabs) restriction enzyme. Eighty μL of 10% SDS were added and samples were then incubated at 65 °C for 30 minutes to inactivate the DpnII enzyme. Samples were brought up to 7 mL and proximity-ligated using 4000 units of T4 DNA ligase (NEB) overnight at 16 °C. Crosslinks were reversed by addition of 30 μL of proteinase K and incubation overnight at 65 °C with shaking. Thirty μL of RNase A were then added and samples were incubated at 37 °C to eliminate RNA. Samples were phenol/chloroform/isoamyl alcohol (Sigma)-extracted and DNA was ethanol precipitated. A subsequent restriction digestion was performed in 500 μL total volume using 50 units of Csp6I (Fermentas). Samples were incubated at 65 °C for 30 minutes to inactivate the enzyme and another proximity ligation reaction was performed in 14 mL total volume using 6000 units of T4 DNA ligase (NEB). DNA was again phenol/chloroform-extracted and ethanol precipitated. DNA was resuspended in 10 mM Tris pH 8.0 and purified using QIAquick PCR purification kits. DNA was quantified by Qubit fluorometry. DpnII/Csp6I 4C templates were PCR-amplified using Expand Long Template PCR System (Roche). In brief, 400 ng of template was used as input in 50 μL PCR reactions using single barcode primers containing the *SOX2* promoter bait, as listed in Table 3. Libraries were isolated using QIAquick PCR purification (Qiagen) and quantified by Qubit and KAPA qPCR (Roche) prior to sequencing.

**Table 3.**
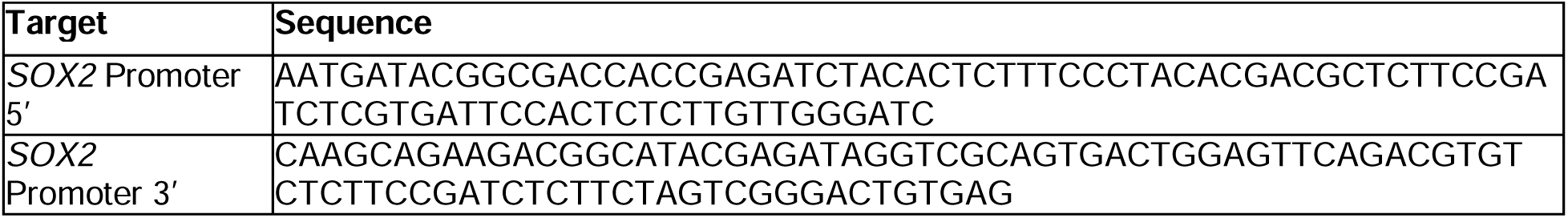
4C-Sequencing Primers: Sequences containing 4C bait sequences and sequencing adapters.

### Preparation of single cell RNA sequencing libraries from teratoma tissues

Previously harvested, flash-frozen teratoma tissue samples were lysed by addition of 0.1 mL of freshly prepared 0.1X lysis buffer (10 mM Tris-HCl, 10 mM NaCl, 3 mM MgCl_2_, 0.1% Tween-20, 0.1% NP40, 0.01% digitonin, 1% BSA, 1 mM DTT, and 1 U/mL Sigma protector RNase inhibitor), tissue homogenized with microfuge tube pestles, and incubated on ice for 10 minutes. One mL of chilled wash buffer (10 mM Tris-HCl pH 7.4, 10 mM NaCl, 3 mM MgCl_2_, 1% BSA, 0.1% Tween-20, 1 mM DTT, and 1 U/mL of RNase inhibitor) was added and the mixture was strained through 70 and 40 µm filters. The resulting solution of nuclei and debris was stained with 5-AAD and the mixture sorted for 5-AAD-positive nuclei on a FACSAria cell sorter. Resulting nuclei were immediately processed to generate single cell RNA-sequencing libraries using the 10X Genomics protocols in the Single Cell 3’ Reagent Kits v2 User Guide. Data were processed, normalized, integrated for all specimens, and clustered together using Seurat packages based on RNA. Only cells with greater than 750 features and fewer than 0.10% mitochondrial genes were included in the analysis. Initial labeling of all cells was performed using the reference mapping feature of Seurat against data from McDonald and colleagues.^52,85^

### Next-generation sequencing

Libraries made according to preceding ChIP, 4C, and HiC library protocols were quantified by Qubit (Thermo Fisher) or KAPA qRT-PCR (Roche) and library quality assessed by Tapestation High Sensitivity (Agilent) electrophoresis. Libraries were pooled as appropriate and sequenced on Illumina Novaseq6000 to achieve the targeted sequencing depth of ∼50 M reads per sample for H3K27me3, ∼30 M reads for H3K27ac, ∼25 M for ATAC-Seq, ∼30 M for RNA-sequencing, ∼5 M reads for 4C libraries, and ∼500 M reads for HiC libraries.

### Processing of sequencing datasets

ChIP-Sequencing and ATAC-sequencing demultiplexed fastq files were trimmed using Trimmomatic, aligned against the Hg19 genome assembly using Bowtie2, and processed to BigWigs and Bed files for visualization using Samtools and Deeptools. Reads Per Genomic Content (RPGC)-normalized files were visualized using the UCSC genome browser or IGV (hg19 chr3:181000000-182500000) with the exception of H3K27ac in rosette NSCs, which was processed using MACS2 to produce Poisson p-value adjusted BigWigs.^84,86^ Peak calling and differential peak analysis was performed using MACS2. HiC datasets were processed using the HiC-bench suite.^87^ 4C-sequencing was mapped against a fragment library of the DpnII-Csp6I digested genome using 4Cker.^88^

### Virtual 4C

To investigate locus-specific 3D chromatin interactions from HiC datasets with virtual 4C, we used the Juicer pipeline and 5-10 kb bins. Viewpoints were defined by genomic coordinates of loci of interest as per the hg19 assembly of the human genome.

### qRT-PCR

Quantitative Real Time PCR for gene expression profiling was performed with the Cells-to-Ct kit (Invitrogen), Taqman Gene Expression kit (Invitrogen) and the StepOne Real Time PCR System (Invitrogen) as recommended by the manufacturers. Following the completion of the qPCR, extracted Ct values from the StepOne Software (v2.1) was then assessed to calculate the normalized target gene expression level using the ΔΔCt method to determine relative fold-change. The following TaqMan (Invitrogen) primers/probes were used: *RBFOX3* (Hs01355808), *TUBB3* (Hs00801390_s1), *SLC1A2* (Hs01102423), *SOX2* (Hs01053049_s1), *GAPDH* (Hs02758991_g1), *FOXG1* (Hs01850784_s1).

### FISH (fluorescent *in situ* hybridization) probe synthesis

Oligo probes targeting the *HS1332* and anchor regions within the *SOX2* TAD were designed using paintSHOP^89^ and ordered from IDT as oPools (probe sequences are listed in Supplementary Table 1). Single-stranded DNA (ssDNA) probe pools were diluted to 0.07 ng/μL and amplified using KAPA HiFi HotStart ReadyMix (Roche) with 100 μM forward and reverse primers specific to each probe set. PCR was performed with the following conditions: 95 °C for 3 min; 20 cycles of 98 °C for 20 s, 56 °C for 15 s, and 72 °C for 30 s; then 72 °C for 30 s. Debubbling mix was then added and the same PCR program was repeated twice. PCR products were purified using the QIAquick PCR Purification Kit (Qiagen). Purified double-stranded DNA templates (≥ 1 μg) were transcribed using the HiScribe T7 High Yield RNA Synthesis Kit (NEB) at 37 °C for 4–16 h. RNA was purified using the NucleoSpin RNA Clean-up Kit (Macherey-Nagel). Complementary DNA (cDNA) was then synthesized using SuperScript IV Reverse Transcriptase (Thermo Fisher) with 52 μg RNA template, 1 mM directly conjugated, fluorescently labeled RT primer, and 10 mM dNTPs (NEB, Cat. No. N0447S). Probes targeting *HS1332* received RT primers labeled with Cy5 (Cytiva), while those targeting the anchor control region used those labeled with Alexa fluor 555 (Thermo). Reactions were incubated at 50 °C for 4 h. Unincorporated primers were digested with Exonuclease I (NEB) at 37 °C for 15 min, followed by heat inactivation in 0.5 M EDTA, pH 8.0, at 80 °C for 20 min to generate RNA:DNA hybrids. RNA:DNA hybrids were purified using the Zymo Quick-RNA Miniprep Kit (Zymo Research, Cat. No. R1054). Residual RNA was digested using RNase H (NEB) and RNase A (Thermo Scientific) sequentially (37 °C for 2 h; 70 °C for 20 min; 50 °C for 60 min). The resulting single-stranded DNA (ssDNA) probes were purified using the Zymo Quick-RNA Miniprep Kit and eluted in 100 μL nuclease-free H_2_0.

The FISH probe library targeting a control locus (chr11, *HBB* locus) was ordered from Arbor Biosciences.

### FISH sample staining

We performed DNA FISH based on previously published protocols with some modifications.^90–92^ All steps were performed at RT unless otherwise specified. Samples were prepared on 1.5H coverslips, fixed in 4% PFA/PBS, rinsed three times in PBS and processed on the same day in 6-well plates. They were then permeabilized with 0.5% Triton X-100 in PBS for 10 min and rinsed for 3 min with PBS twice. Cells were incubated for 5 min in 0.1 N HCL, and rinsed three times for 1 min in 2XSSC (saline sodium citrate), then three times for 3 min in PBS. Cells were incubated in RNAse A at 37 °C for 1 h in a humid environment. Samples were pre-hybridized in preheated Hybridization #1 buffer [0.1% (v/v) Tween-20, 50% (v/v) formamide, 2XSSC] at 37 °C for 30 min in a humid environment. Meanwhile, the probe mix was prepared by complementing 2 pmol of each probe set with Hybridization #2 buffer [0.1% (v/v) Tween-20, 50% (v/v) formamide, 10% (v/v) dextran sulfate, 2XSSC] up to a 10 mL volume. The probe mix was then denatured for 5 min at 85 °C in the dark and chilled for 5 min on ice. Twenty mL of probe mix were added to each coverslip, which was then sealed onto a microscope slide with rubber cement in dark for 5 mins.

Sample chromatin was denatured by placing the slides for 5 mins at 85°C on an inverted heat block submerged in an 85 °C water bath. Slides were then transferred to a humidified slide tray and incubated overnight in the dark at 42 °C. The next day, the rubber cement was peeled off and the coverslip floated off the slides in a 2XSSC reservoir. Coverslips were then transferred to 6-well plates pre-filled with probe wash buffer [0.1% (v/v) Tween-20, 50% (v/v) formamide, 5XSSC] and were incubated at 42 °C for 50 min in a humidified environment. This step was repeated with fresh probe wash buffer, then twice for 5 min in 5XSSCT (5XSSC with 0.1% Tween-20), and 5 min in PBS. Cells were stained with 4′,6-diamidino-2- phenylindole (DAPI) for 5 min and rinsed in PBS. Coverslips were mounted on Prolong Gold, allowed to cure for 24 h in the dark, sealed with nail polish, and imaged.

### FISH sample imaging and analysis

FISH imaging was performed on a Nikon Eclipse Ti-2 inverted epifluorescence microscope controlled by MicroManager,^93^ equipped with a Prime BSI Express camera, using a 100x oil-immersion lens (NA=1.4). Image stacks were captured with voxel size 64.5 x 64.5 x 250 nm. Slides were imaged using four channels: 405 nm (DAPI), 488 nm (chr11 control probe targeting the *HBB* locus), 532 nm (Alexa Fluor 555; chr3 anchor), and 637 nm (Cy5; *HS1332* sequence) excitation lasers.

Nuclei were segmented in 2D from maximum intensity projections of the DAPI signal using Cellpose.^94^ FISH spots were detected in each channel using AIRLOCALIZE,^95^ and the FISH spots were individually mapped to their matching nuclei based on their x,y coordinates using a custom MATLAB script.

### Flow cytometry

Human ESC colonies were dissociated using TrypLE (Gibco) and 1x10^6^ cells were centrifuged at 300xg for 5 min before being washed with Dulbecco’s phosphate buffered saline (Gibco). Cell pellets were then resuspended in fluorescence activated cell sorting (FACS) buffer (0.5% BSA in DPBS with 2 mM EDTA). Each sample was incubated with 2 μL of SSEA4 488 (Miltenyi), TRA-1-81 PE (Miltenyi) or an IgG1 isotype control antibody with matching fluorophore (Miltenyi) at 4 °C for 10 min. Cells were pelleted at 300xg for 5 min and washed twice with FACS buffer, before being resuspended in buffer with 100 ng DAPI for dead cell exclusion. Samples were analyzed by flow cytometry (BD LSRFortessa II – BD Biosciences) and resulting data was analyzed using FlowJo™ v10.8 Software (BD Life Sciences).

Intrinsic GFP expression from the *Hes5::GFP* reporter was assessed by flow cytometry by pelleting 1x10^6^ hESCs or rosette NSCs per condition, washing twice with DBPS, and resuspending in FACS buffer containing 100 ng of DAPI for dead cell exclusion. Samples were run and analyzed as described above.

### Whole genome sequencing and pseudokaryotype generation

Genomic DNA was extracted from freshly harvested 5x10^6^ hESCs per condition using the DNeasy (Qiagen) DNA collection kit according to the manufacturer’s protocol. Whole-genome libraries were prepared by the NYU Genome Technology Center using the Illumina DNA Prep PCR-Free Library Kit and sequenced on an Illumina NovaSeq X with a target of 50X genomic coverage. Demultiplexed fastq files were trimmed of adapters and filtered of low-quality reads using Trim Galore with Phred quality scores below 20. Reads were aligned against the UCSC HS1 telomere-to-telomere human genome using Bowtie2 to generate BAM files, and duplicates were removed using picard tools. The resulting BAM files were used to create BigWig files utilizing deeptools with 500 kb binning for purposes of pseudokaryotypes or 10 bp binning for CRISPR target deletion confirmation.^83^ Pseudokaryotypes were generated from BigWig files using Karyoploter.^96^

### Western blot

10 million cells per condition were lysed in RIPA buffer (Thermo Fisher) supplemented with Halt protease inhibitor cocktail (Thermo Fisher). Samples were incubated on ice for 10 min and then sonicated in a water-bath Bioruptor (Diagenode) at medium power setting for 10 cycles of 15 s “On” and 60 s “Off” at 4 °C. Lysates were centrifuged at 15,000xg for 10 min at 4 °C. Protein concentrations were determined using the DC protein assay kit II (Bio-Rad). Laemmli buffer (Bio-Rad) containing β-mercaptoethanol was then added to protein lysates. Samples were incubated at 95 °C for 5 min. Twenty μg of protein per sample were loaded onto SDS-PAGE gels for separation and transferred to 0.2 μm nitrocellulose membranes (Bio-Rad). After blocking the membranes in 2% bovine serum albumin (BSA) in TBS-Tween for 1 h at RT, they were simultaneously incubated with antibodies against SOX2 (R&D Systems) or β- Actin at 1:200 and 1:1000, respectively. Blots were incubated at 4 °C overnight and visualized with two simultaneous fluorescent Alexa Fluor Plus–conjugated secondary antibodies. Images were acquired using the iBright FL1000 system (Invitrogen). Densitometric quantification of band intensities was conducted in ImageJ.

### Immunofluorescence microscopy

Cell cultures were fixed with 4% paraformaldehyde (PFA) for 30 min at RT, followed by block and permeabilization with 10% bovine serum albumin (BSA) in PBS supplemented with 0.1% Triton X-100 for 1 h at RT. Cells were then incubated with primary antibodies (Table 4) in 1% BSA in PBS + 0.1% Triton X-100 at 4°C overnight. The next day, cells were washed with PBS + 0.1% Triton X-100 and stained with donkey AlexaPlus IgG (H+L) secondary antibodies for 1 h at RT. Nuclei were counterstained with 500 ng/mL 4′,6-diamidino-2-phenylindole (DAPI) for 10 min at RT. Microscopy was conducted on a Zeiss Axio-observer microscope.

**Table 4.**
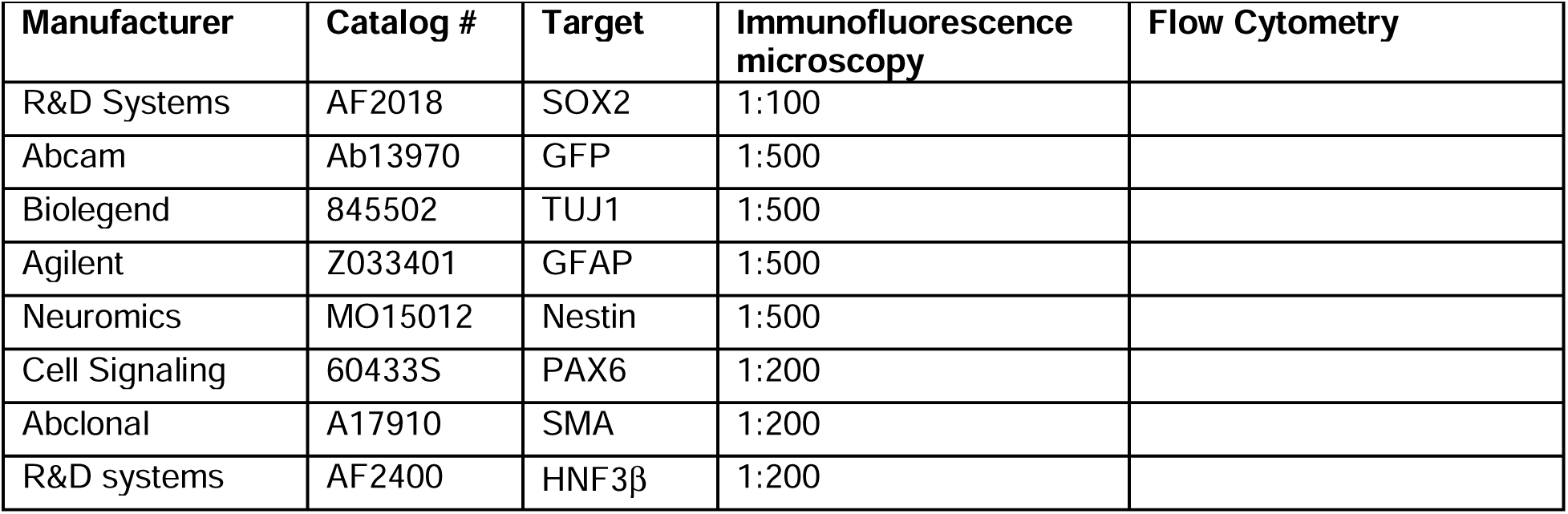
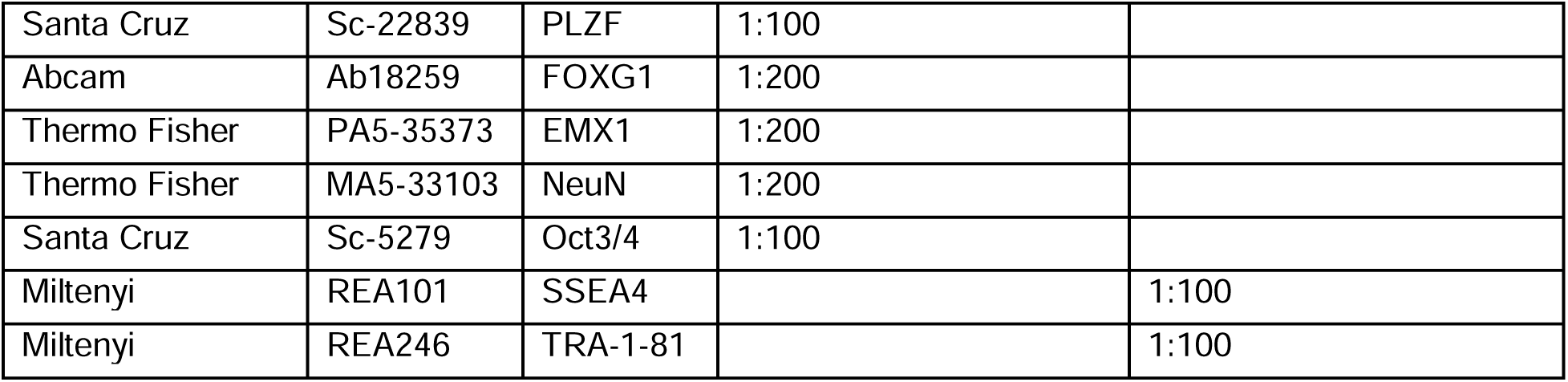
Antibodies and Dilutions Used.

| Manufacturer | Catalog # | Target | Immunofluorescence microscopy | Flow Cytometry |
| --- | --- | --- | --- | --- |
| R&D Systems | AF2018 | SOX2 | 1:100 |  |
| Abcam | Ab13970 | GFP | 1:500 |  |
| Biolegend | 845502 | TUJ1 | 1:500 |  |
| Agilent | Z033401 | GFAP | 1:500 |  |
| Neuromics | MO15012 | Nestin | 1:500 |  |
| Cell Signaling | 60433S | PAX6 | 1:200 |  |
| Abclonal | A17910 | SMA | 1:200 |  |
| R&D systems | AF2400 | HNF3β | 1:200 | |
| Santa Cruz | Sc-22839 | PLZF | 1:100 |  |
| Abcam | Ab18259 | FOXG1 | 1:200 |  |
| Thermo Fisher | PA5-35373 | EMX1 | 1:200 |  |
| Thermo Fisher | MA5-33103 | NeuN | 1:200 |  |
| Santa Cruz | Sc-5279 | Oct3/4 | 1:100 |  |
| Miltenyi | REA101 | SSEA4 |  | 1:100 |
| Miltenyi | REA246 | TRA-1-81 |  | 1:100 |

### Teratoma tissue harvesting, sectioning, and staining

Each teratoma was partially dissected to obtain small samples for flash freezing and cryopreservation for downstream analysis by sequencing assays. The remaining teratoma tissue was immediately transferred to 4% PFA and incubated at 4 °C for 24 hours. Samples were then transferred to 20% sucrose to dehydrate overnight at 4 °C. After tissue dehydration was complete, samples were embedded into optimal cutting temperature (OCT) medium, mounted onto cutting blocks, and cryosectioned at 20 μm thickness onto glass slides. Slides were immediately transferred to -20°C for storage until immunostaining, which was performed as described above, or H&E staining. The presence or absence of neural rosettes was scored by pathologists blinded to experimental groups.

### Tissue clearing and immunostaining of cerebral organoids

Cerebral organoids were cleared and immunostained as previously published.^97^ In brief, organoids were fixed in 4% PFA and then incubated in PBS with 0.1% Tween-20. Organoids were washed with organoid wash buffer (PBS, 0.1% Triton X-100, 0.2% BSA) and blocked at RT. They were then incubated with primary antibodies (Table 4) for 24 hours at 4°C. Following primary antibody incubation, organoids were washed 3 times for 2 hours per wash. They were then incubated with secondary antibodies [donkey AlexaPlus IgG (H+L) diluted to 1:500 in organoid wash buffer] for 24 hours at 4 °C. After 3 washes with organoid wash buffer (2 hours per wash), they were stained with 500 ng/mL DAPI. Organoids were then gently centrifuged at 70xg for 3 minutes and resuspended in 150 μL of FocusClear (CelExplorer Labs) for 1 hour or until visually clarified. Organoids were then mounted in 40 μL of FocusClear on glass slides with glass #1 coverslips (Fisherbrand) and imaged on a Zeiss LSM800 confocal microscope.

### Quantification and statistical analysis

Statistical analysis was performed using GraphPad Prism (version 8.4.3). Population statistics are represented as mean ± standard error of the mean (SEM) throughout. Statistical significance was calculated using Student’s t-test, one-way analysis of variance (ANOVA) or two-way ANOVA with Tukey’s and Sidak’s *post hoc* test for multiple comparisons, χ2 for trend and Benjamini-Hochberg multiple testing procedure. P values <0.05 were considered statistically significant (*p<0.05; **p<0.01; ***p<0.001; ****p<0.0001). For GO term statistical analysis of scRNA-seq data, differential expression (DE) testing was based on the non-parametric Wilcoxon rank sum test with multiple comparison correction by FDR (False Discovery Rate). For gene expression analysis of scRNA-seq data, statistical difference among groups was determined by nonparametric Kruskal-Wallis test with pairwise Wilcoxon *post hoc* test corrected by Benjamini-Hochberg FDR for multiple comparisons, or Mann Whitney U test with Benjamini-Hochberg FDR correction.

## KEY RESOURCES

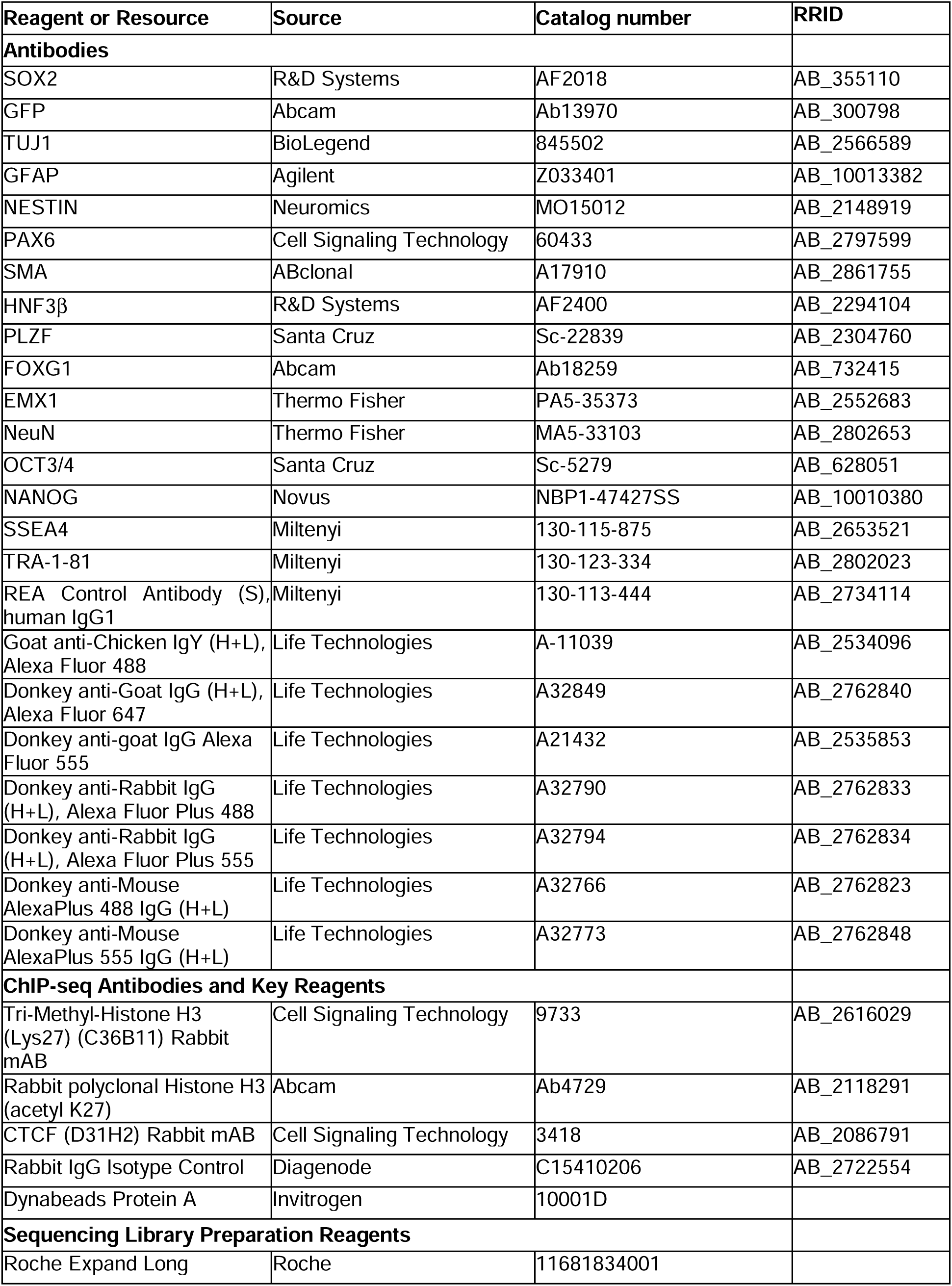
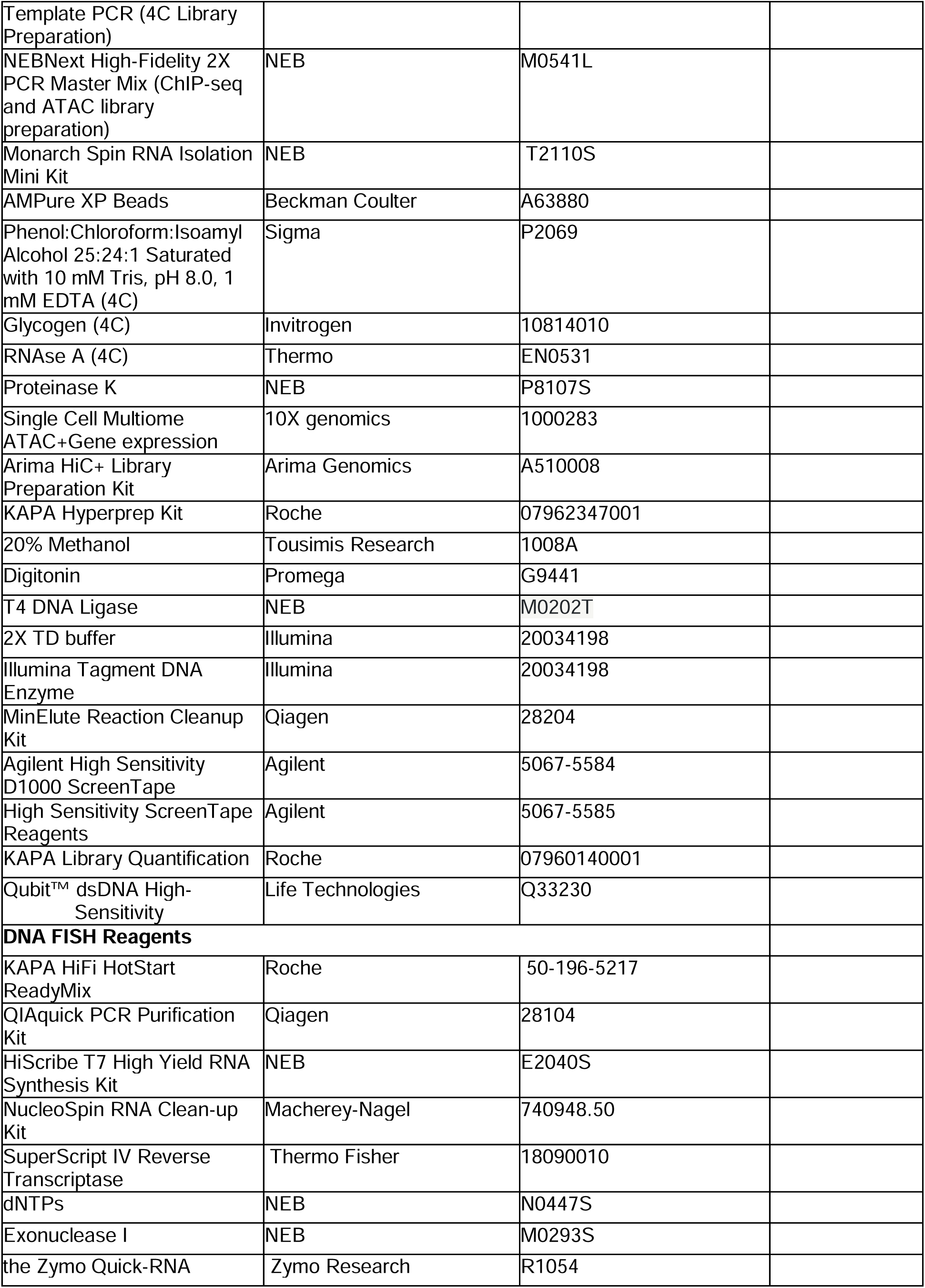
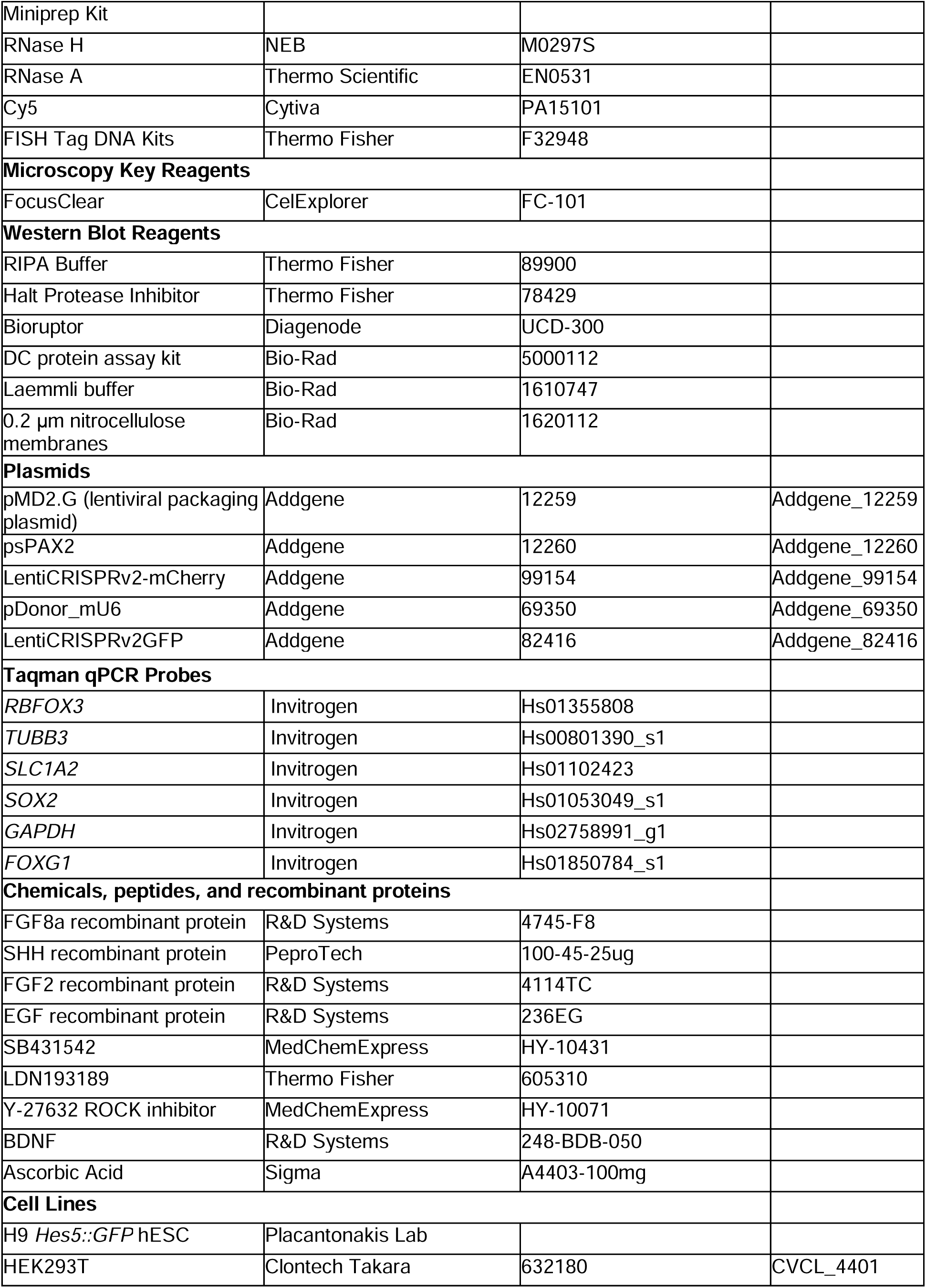
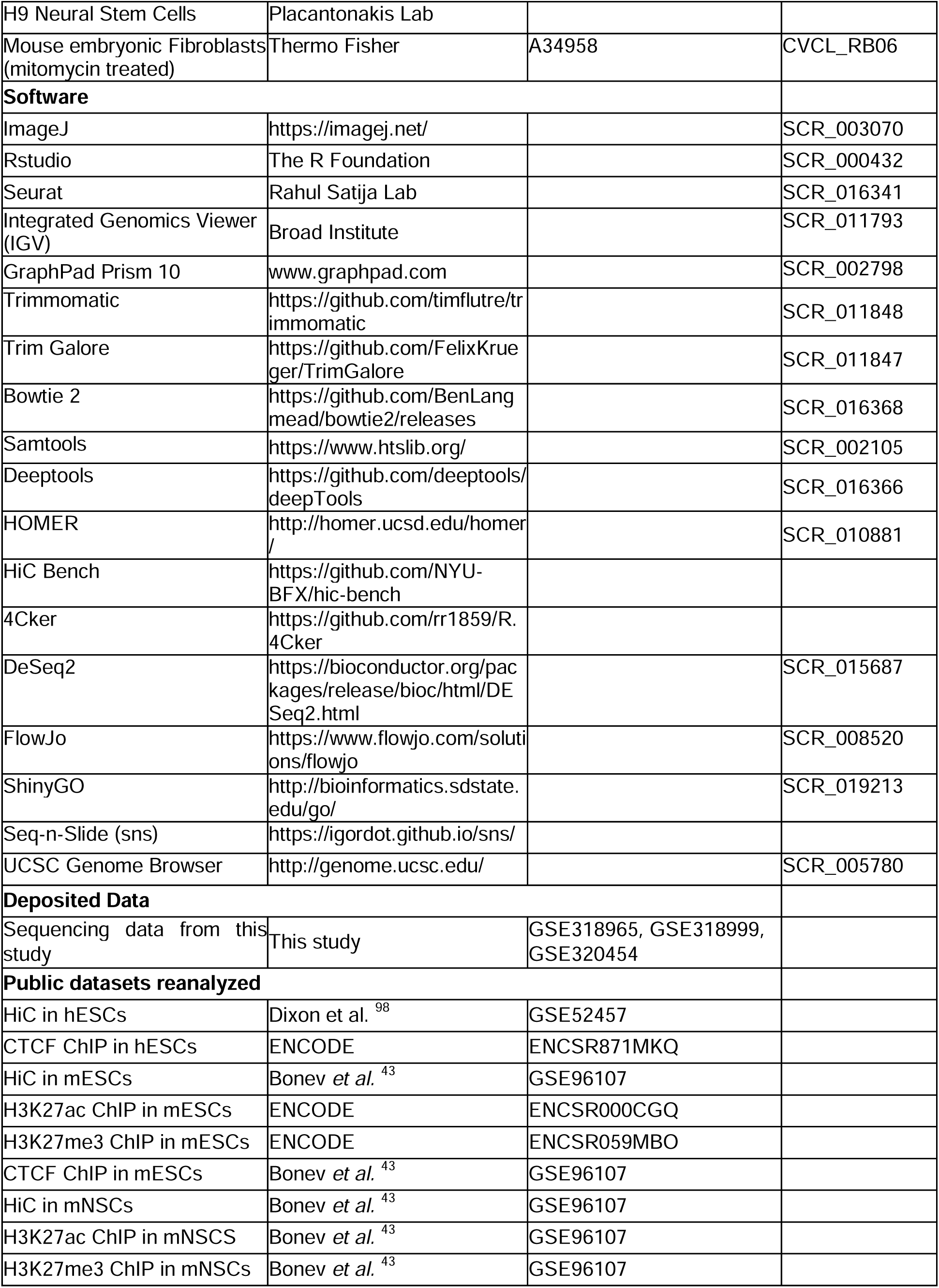
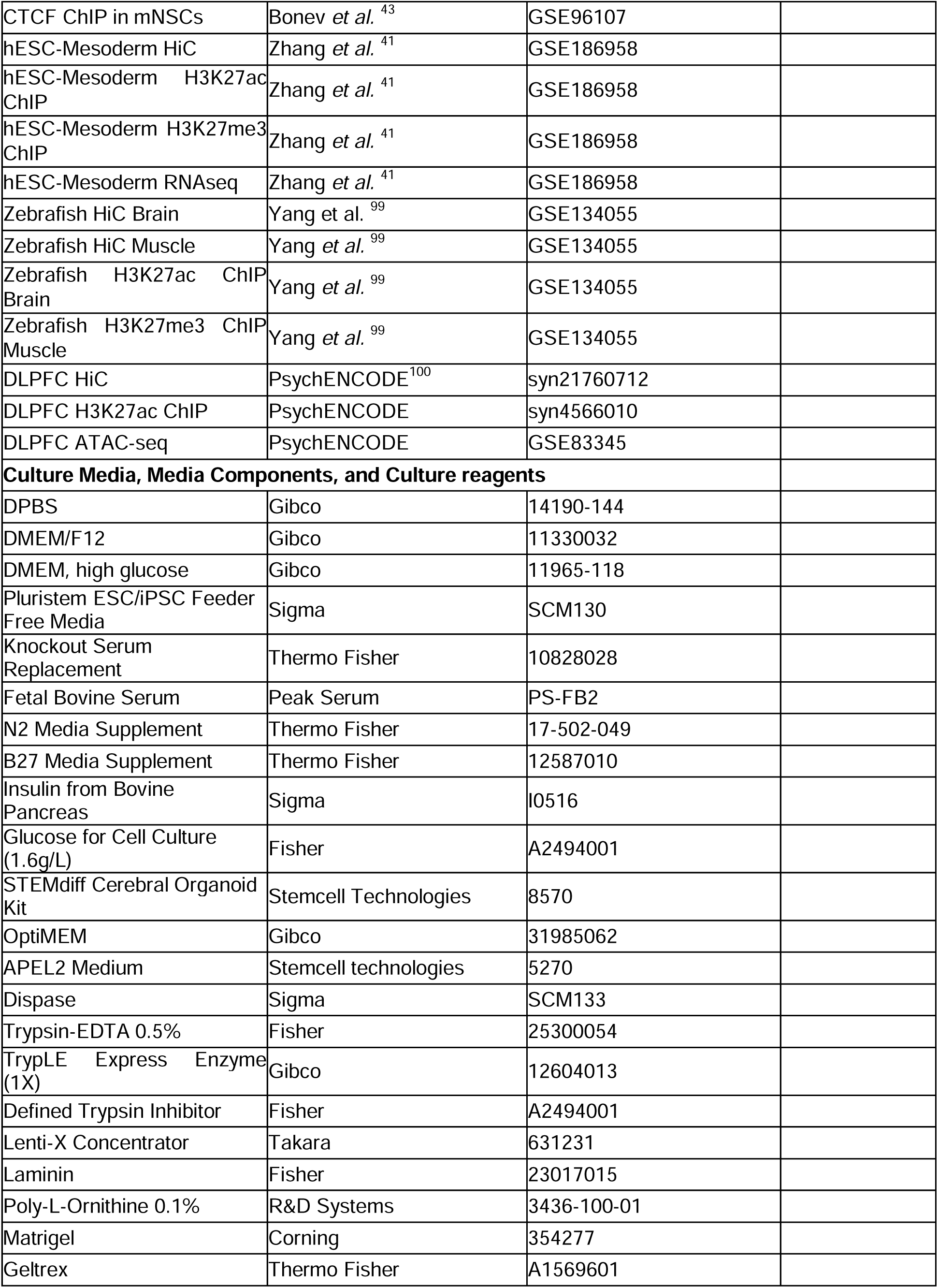
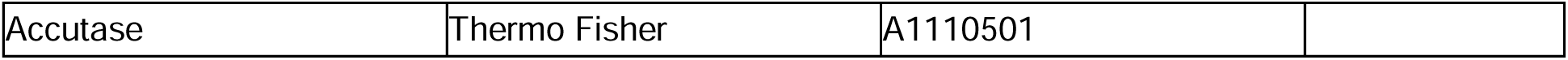

