## Supplemental figure legends for "A developmentally regulated long-range enhancer-promoter contact mediates human neural development"

### Figure S1. 3D chromatin organization and chromatin modifications within the SOX2 TAD in distinct developmental stages and tissues

(A) Genome browser view of the *SOX2* TAD showing ATAC-sequencing in hESCs and NSCs, and H3K27ac ChIP-sequencing done in NSCs, hESCs, and mesodermal progenitors (GSE186958).
(B) Genome browser view of the *SOX2* TAD with HiC, ChIP-sequencing for H3K27ac, HeK27me3 and CTCF, RNA-sequencing, and virtual 4C-sequencing (inferred from HiC data) in hESCs (our own data), hESC-derived mesodermal progenitors (GSE186958), and NSCs (our own data).
(C) Genome browser view of the zebrafish genome containing the *SOX2* locus and enhancer sequence homologous to *HS1332*. HiC and H3K27ac ChIP-sequencing peaks for brain and muscle tissues are displayed (GSE134055).

### Figure S2. Genotyping and Whole Genome Sequencing (WGS) for ΔHS1332 and ΔnMotif hESCs

(A) Gel electrophoresis of genotyping PCR products in control *ROSA26* and modified Δ*HS1332* and Δ*nMotif* hESCs, and Sanger sequencing of the PCR amplicons.
(B) WGS reveals hemizygous deletion of the *HS1332* sequence in Δ*HS1332* hESCs.
(C) WGS demonstrates homozygous deletion of the *nMotif* sequence in Δ*nMotif* hESCs.

### Figure S3. FISH analysis in ΔHS1332 hESCs and rosettes

(A) Visualization of the *SOX2* TAD showing the enhancer and anchor FISH probes in relation to ChIP-sequencing H3K27ac data.
(B) Quantification of the relative number of FISH enhancer spots normalized to the number of FISH anchor spots in hESCs and rosettes in the *ROSA26* vs Δ*HS1332* conditions.
(C) Quantification of the raw number of FISH enhancer spots in hESCs and rosettes in the *ROSA26* vs Δ*HS1332* conditions.
(D) Quantification of the raw number of FISH anchor spots in hESCs and rosettes in the *ROSA26* vs Δ*HS1332* conditions.
(E) Quantification of the raw number of FISH chr11 control spots in hESCs and rosettes in the *ROSA26* vs Δ*HS1332* conditions.

### Figure S4. WGS-inferred pseudokaryotypes in hESCs

Pseudokaryotypes inferred from WGS in *ROSA26*, Δ*HS1332* and ΔnMotif hESCs (500 kb genomic bins). Sequences were aligned against the UCSC HS1 telomere-to-telomere human genome.

### Figure S5. Flow cytometric assessment of pluripotency cell surface markers

(A) Representative flow cytometry histogram plots for SSEA4 and TRA-1-81 in *ROSA26*, Δ*HS1332* and ΔnMotif hESCs.
(B) Percentage of cells positive for each surface marker (n=3 biological replicates, one-way ANOVA p>0.05). ns, not significant.

### Figure S6. H&E staining of teratomas and quantification of repressive chromatin modifications in loci within the SOX2 TAD

(A) Representative H&E-stained tissue from *ROSA26*, Δ*HS1332*, and Δ*nMotif* teratomas.
(B) Blinded reviewer scoring of teratoma from the three experimental conditions for presence or absence of neuroectodermal tissue (χ2 test for trend p=0.0363).

### Figure S7. Expression of SOX2, NOTCH1 and HES/HEY transcripts in neuroectodermal clusters (cycling progenitors, early neurons, radial glia) in teratoma scRNA-seq datasets

(A) Violin plots showing expression of *SOX2*, *NOTCH1*, *HES1*, *HES5*, *HEY1* and *HEY2* in these clusters (Kruskal-Wallis test with pairwise Wilcoxon *post hoc* test corrected by Benjamini-Hochberg FDR for multiple comparisons). ns, not significant.
(B) Box-whisker plots describing expression of each gene only in cells positive for the annotated gene (Mann-Whitney U test with Benjamini-Hochberg FDR correction).

### Figure S8. Enrichment of H3K27me3 deposition in peaks 1-3 across the ROSA26, ΔHS1332 and ΔnMotif conditions. Peak 1 shows H3K27me3 enrichment in ΔHS1332 hESCs, while peaks 2 and 3 manifest similar enrichment in ΔHS1332 rosettes (n = 2 biological replicates).

**Supplemental Table 1. FISH probes**
