## Supplementary figures and images for "A developmentally regulated long-range enhancer-promoter contact mediates human neural development"

### Figure S1

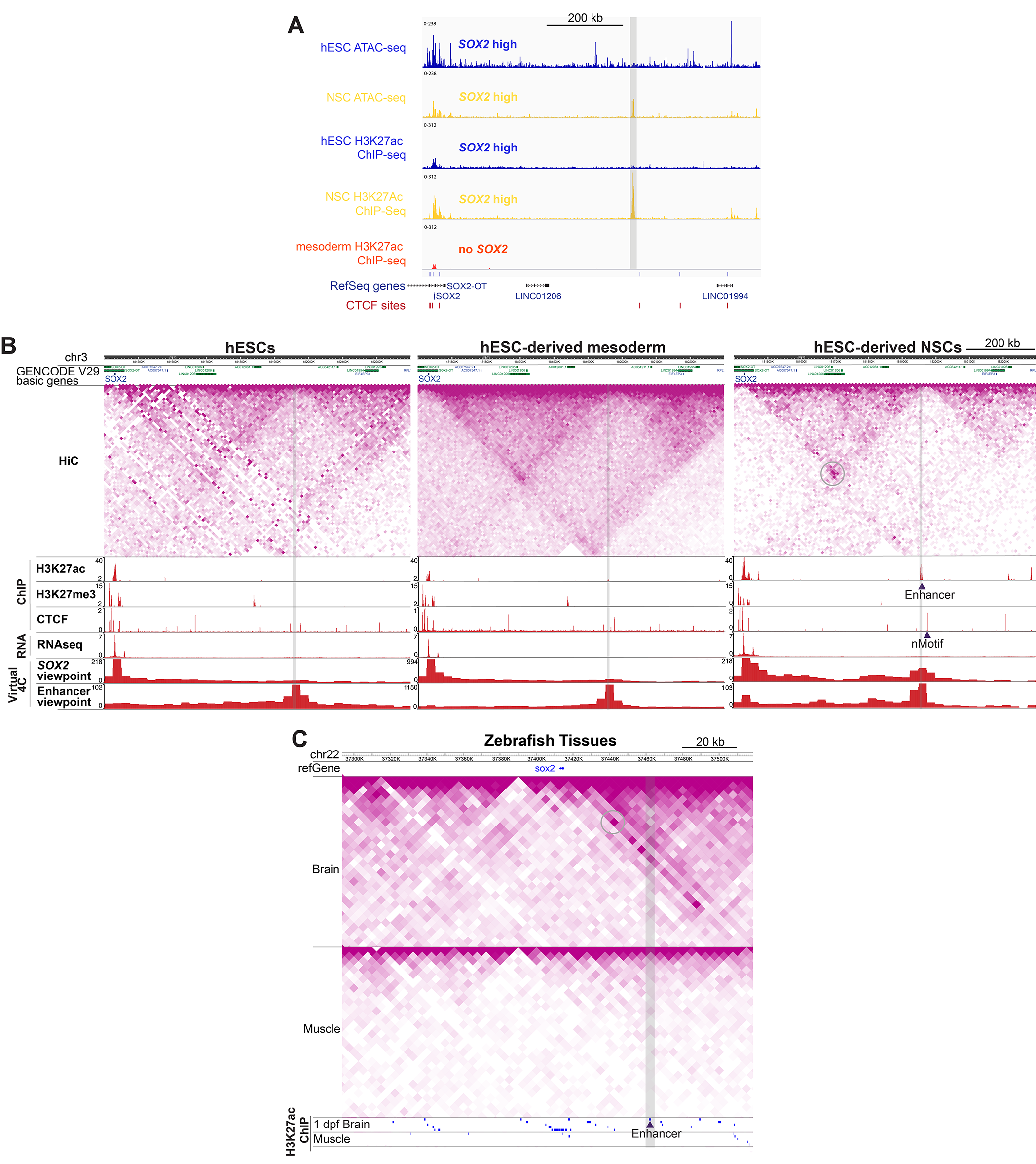

### Figure S2

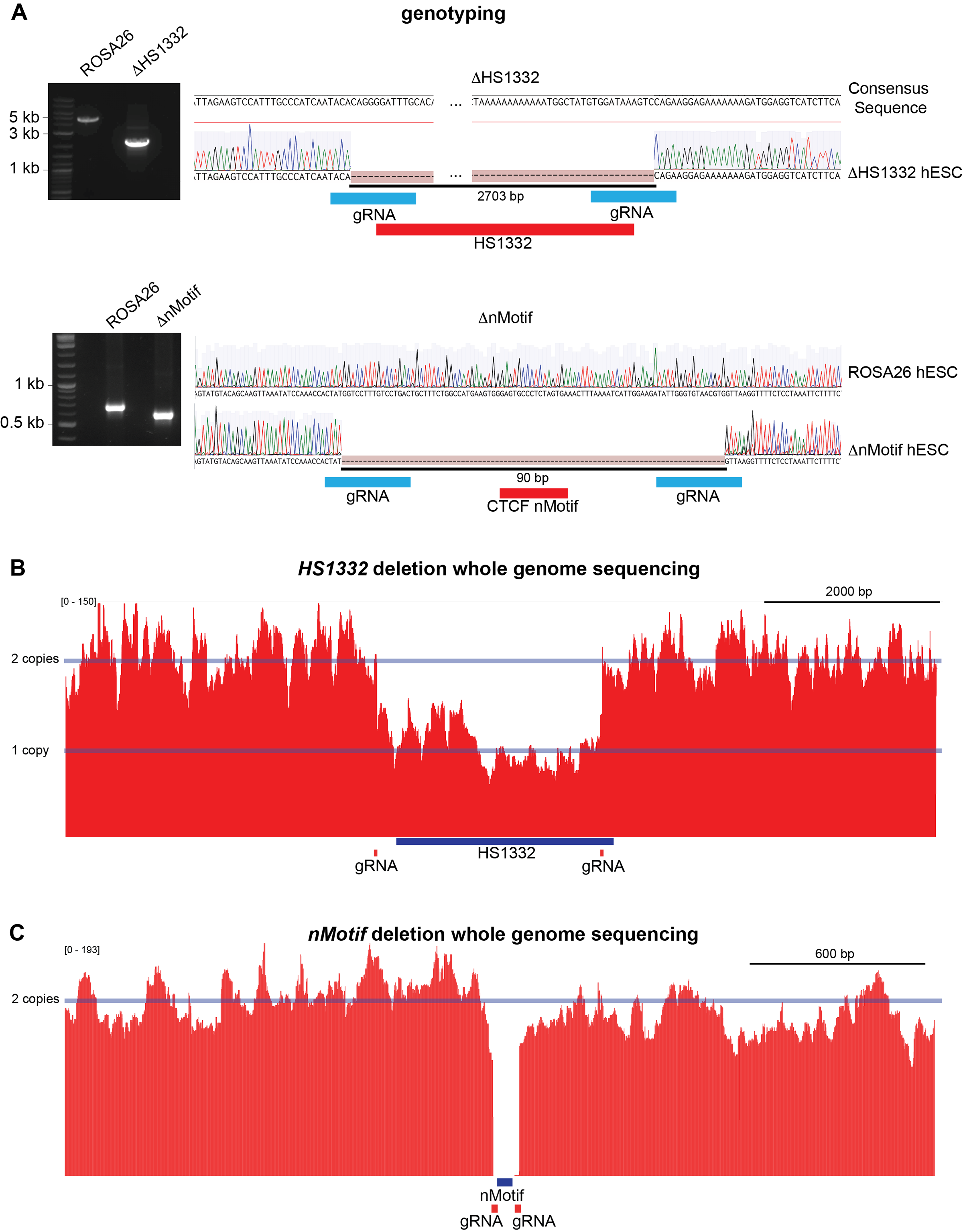

### Figure S3

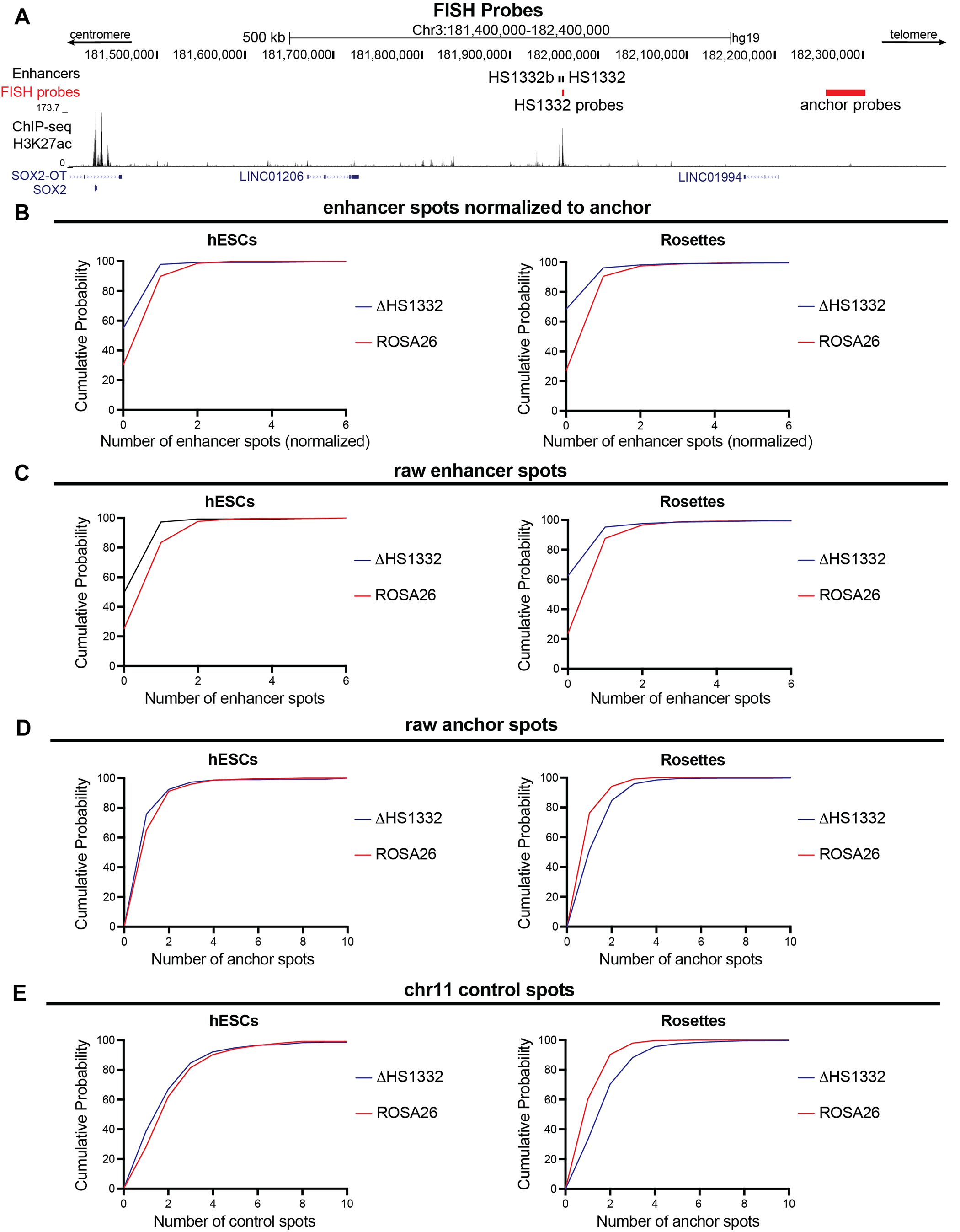

### Figure S4

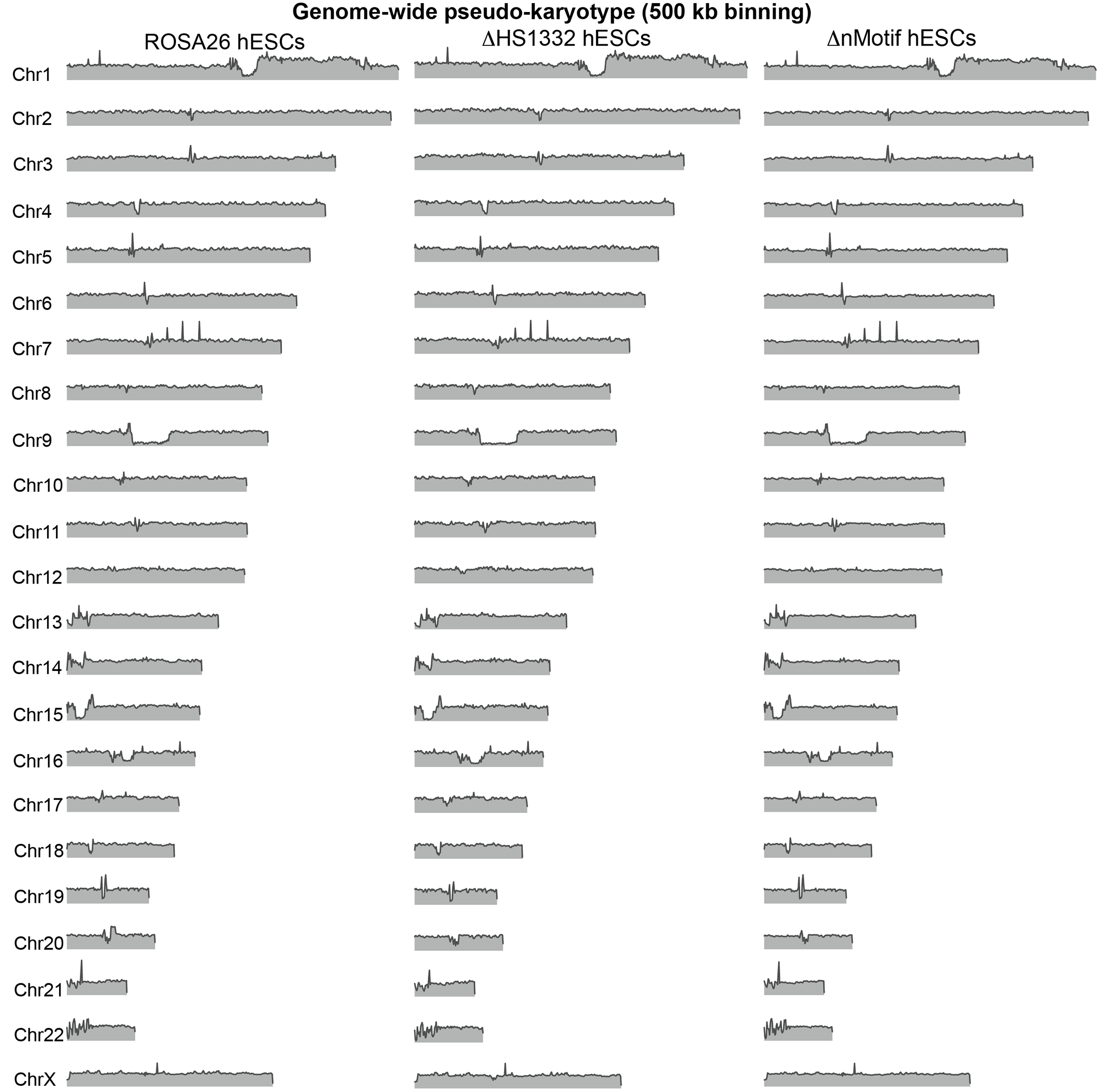

### Figure S5

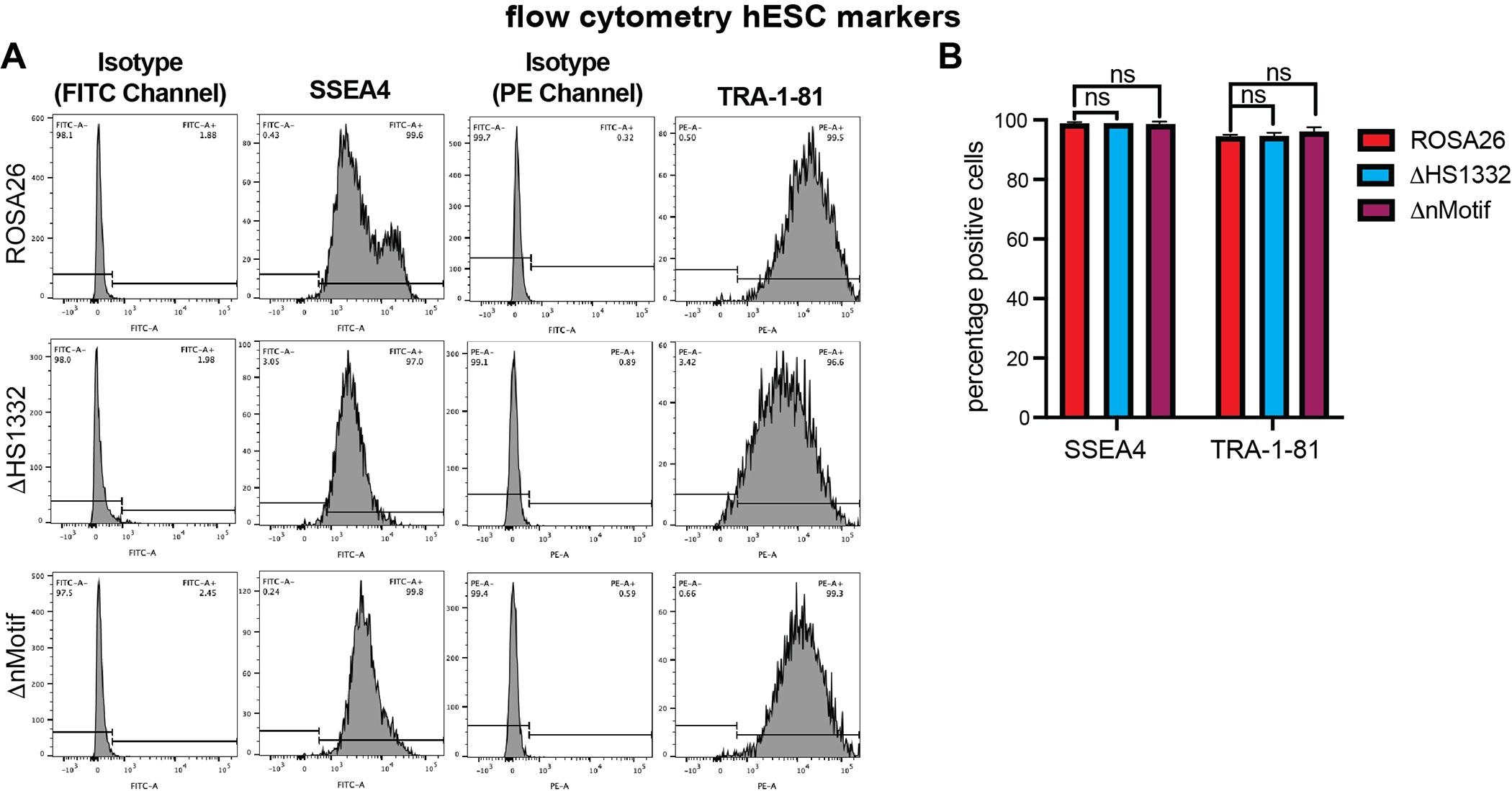

### Figure S6

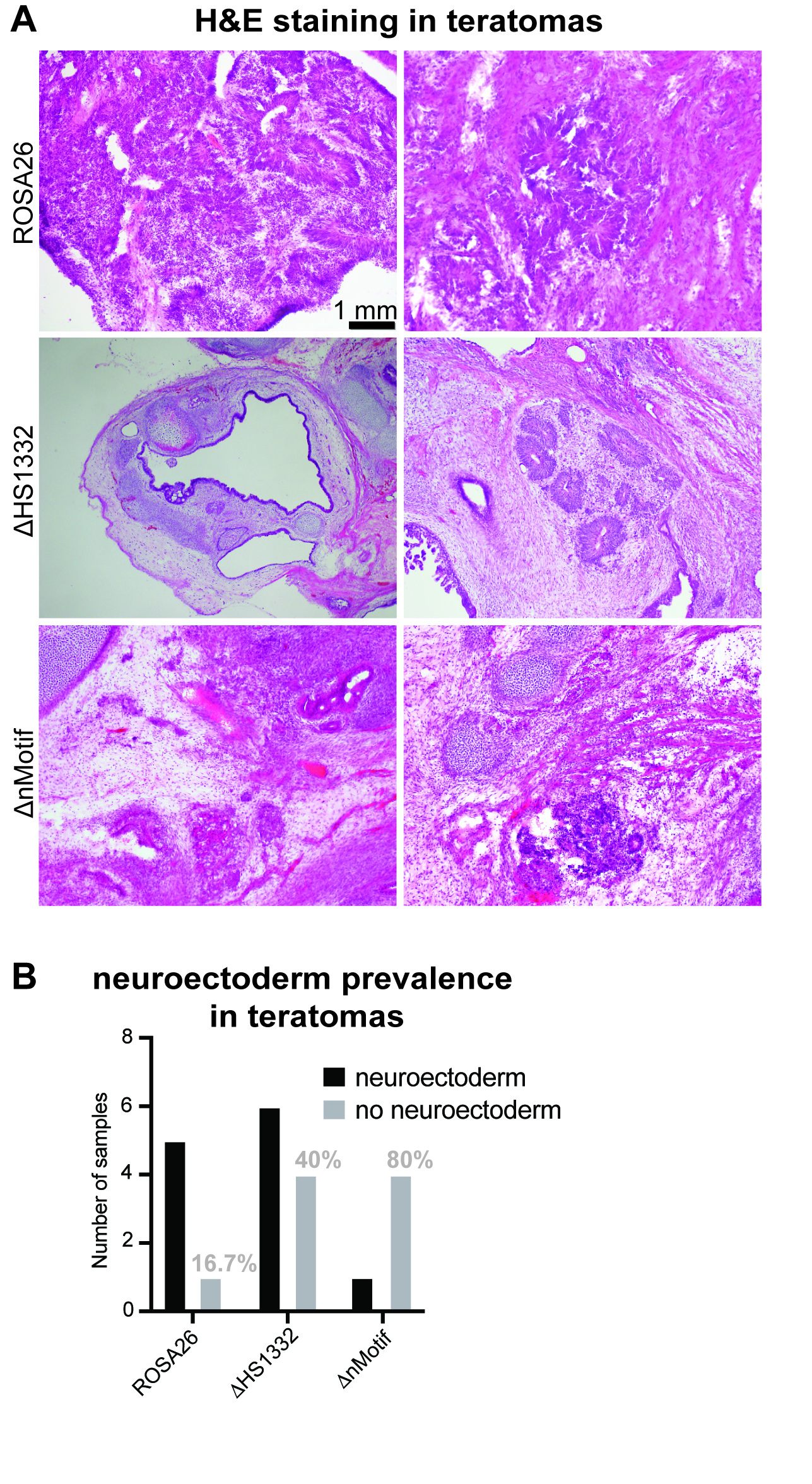

### Figure S7

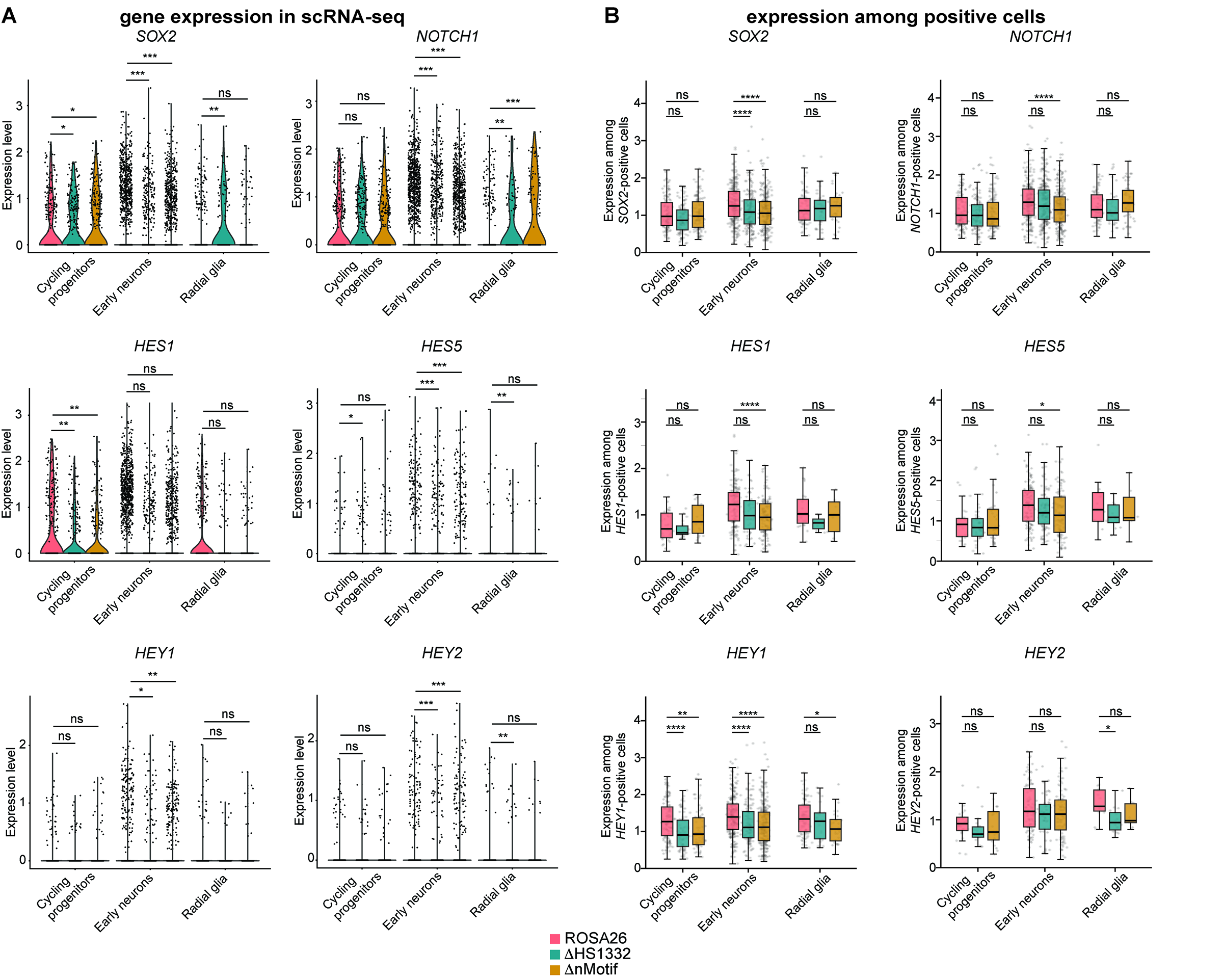

### Figure S8

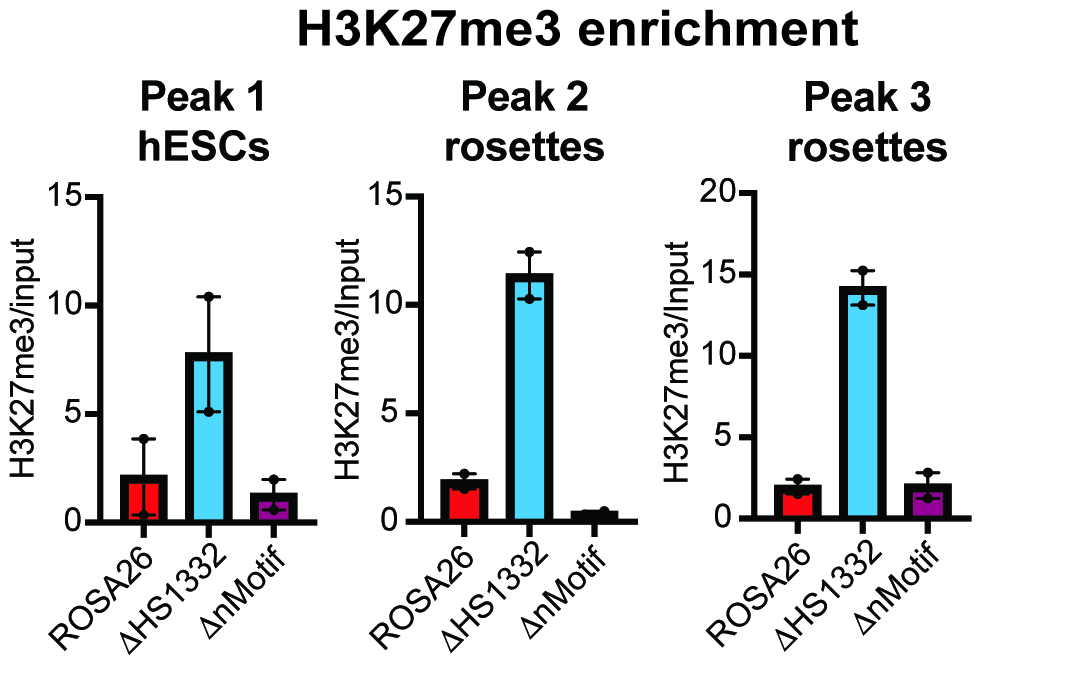
